# TripletGO: Integrating Transcript Expression Profiles with Protein Homology Inferences for High-Accuracy Gene Function Annotations

**DOI:** 10.1101/2021.11.25.470058

**Authors:** Yi-Heng Zhu, Chengxin Zhang, Yan Liu, Gilbert S. Omenn, Peter L. Freddolino, Dong-Jun Yu, Yang Zhang

**Author notes:** Corresponding author(s). (Zhang Y), (Yu D). (Zhu Y), (Zhang C), (Liu Y), (Omenn G), (Freddolino P).

## Abstract

Gene Ontology (GO) has been widely used to annotate functions of genes and gene products. We proposed a new method (TripletGO) to deduce GO terms of protein-coding and non-coding genes, through the integration of four complementary pipelines built on transcript expression profiling, genetic sequence alignment, protein sequence alignment and naïve probability, respectively. TripletGO was tested on a large set of 5,754 genes from 8 species (human, mouse, arabidopsis, rat, fly, budding yeast, fission yeast, and nematoda) and 2,433 proteins with available expression data from the CAFA3 experiment and achieved function annotation accuracy significantly beyond the current state-of-the-art approaches. Detailed analyses show that the major advantage of TripletGO lies in the coupling of a new triplet-network based profiling method with the feature space mapping technique which can accurately recognize function patterns from transcript expressions. Meanwhile, the combination of multiple complementary models, especially those from transcript expression and protein-level alignments, improves the coverage and accuracy of the final GO annotation results. The standalone package and an online server of TripletGO are freely available at https://zhanglab.ccmb.med.umich.edu/TripletGO/.

## Introduction

In the post-genome sequencing era, a major challenge is to annotate the biological functions of all genes and gene products, which are grouped, in the context of the widely used Gene Ontology (GO), into three aspects of molecular function (MF), biological process (BP) and cellular component (CC) [1]. Accurate annotation of gene function provides essential knowledge to disease mechanisms and drug design [2, 3]. Direct determination of the functions of genes via biochemical or genetic experiments is typically time-consuming and laborious, and often incomplete [4]. As a result, a large number of genes in the sequenced genomes have no available function annotation to date. For example, according to official statistics in the neXtProt platform [5], nearly 2,000 protein-coding human genes have yet no known function; for many other organisms of biomedical or industrial importance, annotation rates are substantially lower. To fill the gap between sequence and function, it is urgent to develop efficient computational algorithms for function prediction [6, 7].

Function annotations can be performed at either the protein- or gene-level. In the former case, the function of the query gene is determined by that of its encoded protein, which can be deduced from the protein sequence, structure, or family information [8-18]. However, protein coding genes account for only ∼2% of a typical multicellular eukaryote genome such as that of humans [19]. There are also many genes for non-coding RNAs as well as genes whose coding potential is unknown or ambiguous.

Most gene-level annotation methods deduce GO terms for queries by using a guilt-by-association (GBA) strategy, which is typically based on the similarity of expression profiles between the gene of interest and template genes with known GO annotations [20-22]. The rationale of GBA is reasonable as genes with the same functions often show similar expression profiles. This was supported by the third Critical Assessment of Protein Function Annotation (CAFA) challenge showing that expression profile has a great potential to improve prediction performance [23]. Despite the achievements of current expression profile-based methods, however, challenges remain.

First, it is tricky to define an effective similarity measure of expression profiles as the substitute for functional similarity. In previous work, several unsupervised methods (e.g., Pearson correlation coefficient [24] and mutual rank [25]) and supervised methods (e.g., metric learning for co-expression [21]) have been developed to measure the expression profile similarity in gene function prediction. Unfortunately, these methods cannot achieve optimal performance, because these expression similarity metrics may have no close correlation with functional similarity. Part of the reason is that these methods define the expression similarity in the original space, in which the expression data show a high dimensionality across multiple tissues and complicated distributions; as a result, the measured expression similarity is hardly associated with functional similarity and thus a higher expression similarity (by these metrics) often does not indicate a higher functional similarity. To address this issue, a promising approach is to change the data distribution via feature space mapping [26], in which the expression profiles are mapped from the original feature space to a new embedding space by non-linear functions, and the expression similarity is then associated with functional similarity in this embedding space. The second challenge is that the functional similarity of genes is often difficult to completely capture by one similarity measure. This necessitates the combinations of multiple similarity measures from different biological datasets which may help improve both accuracy and coverage of function predictions [9].

In this work, we proposed and tested a new approach, TripletGO, to integrate multi-source information from both genes and proteins for protein-coding and non-coding gene annotations. First, we extended a supervised triplet network method [27] to assess expression profile similarities in function prediction. In this extended triplet-network pipeline (TNP), the expression profiles are mapped from the original feature space to an embedding feature space via deep neural network learning, where a triplet loss function is designed to enhance the expression profile and gene function correlation. Second, considering that most protein-coding gene functions are performed through proteins and that protein sequence alignments, which are based on 20 amino acids, often provide more specific function associations than nucleotide sequence alignments, we proposed a protein-level method for GO prediction using protein sequence similarity. Finally, a composite model is derived by integrating the output of four complementary GO prediction pipelines, built on the TNP-based expression profiling, genetic sequence alignment, naïve probability, and protein sequence alignment respectively, through an optimal neural network training. TripletGO has been systematically tested on a large set of non-redundant genes collected from eight species, where the results demonstrated significant advantage on accurate GO term prediction over the current state of the art of the field. The standalone package and an online server of TripletGO are freely available through URL https://zhanglab.ccmb.med.umich.edu/TripletGO/.

## Results

### TNP improves expression profile-based GO prediction

TripletGO is a hierarchical approach that takes as input the gene sequence and the transcript expression profile data. Gene ontologies are then created by a set of four complementary pipelines, built on transcript expression profiling, gene sequence alignment, protein sequence alignment and naïve prior statistical calculation respectively, where the final GO models are obtained by a neural network combination (**Figure 1**).

**Figure 1.**
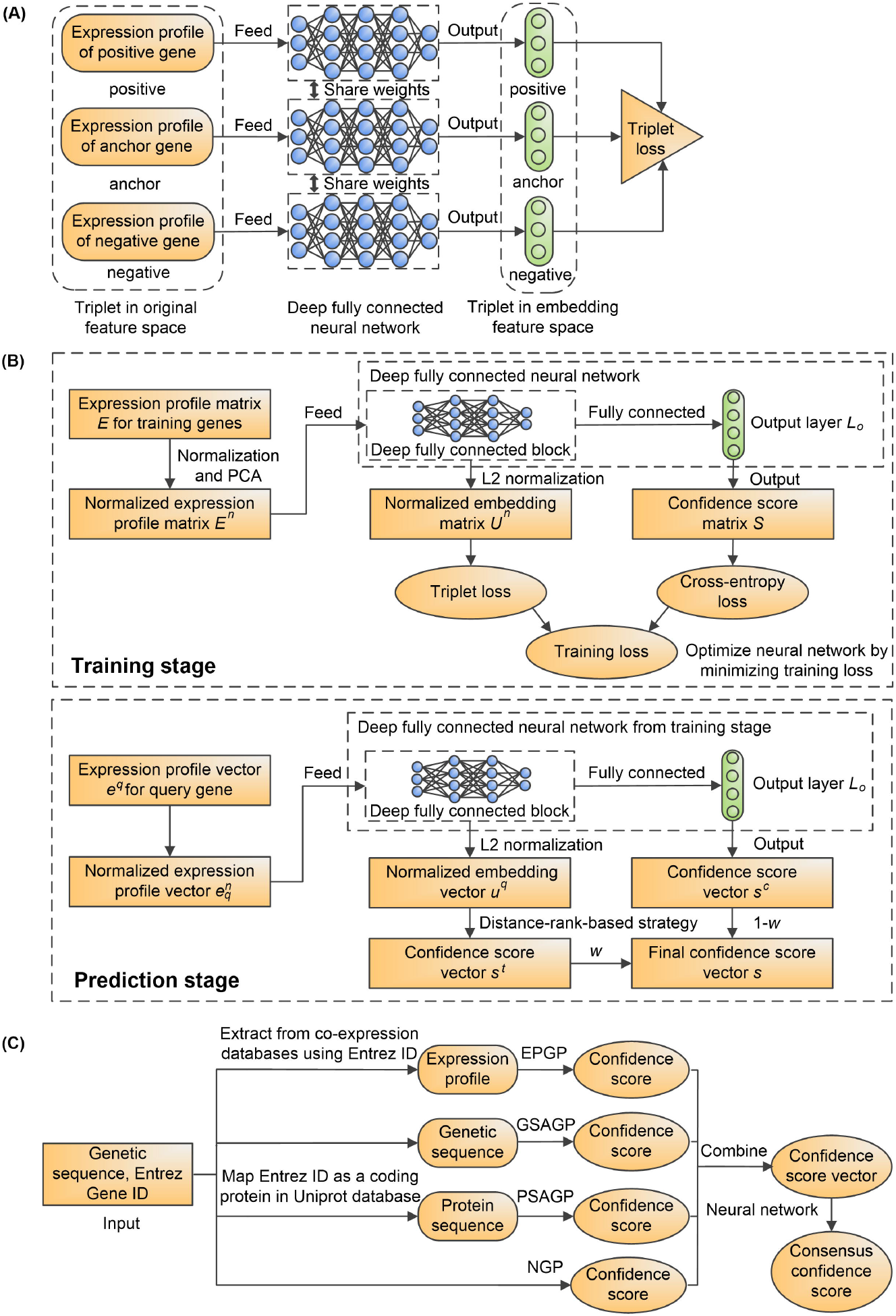
The procedures of TripletGO. **A**. The design of a triplet-network for assessing expression profile similarity. **B**. The triplet network-based pipeline (TNP) for expression profile-based GO prediction. **C**. The flowchart of TripletGO to integrate four complementary pipelines for GO prediction. EPGP: expression profile-based GO prediction, GSAGP: genetic sequence alignment-based GO prediction, PSAGP: protein sequence alignment-based GO prediction, NGP: naïve-based GO prediction.

As TripletGO is centralized by the transcript expression profiling through TNP (**Figure 1B**), we first compare the TNP with five existing methods in expression profile-based GO prediction. These include four unsupervised scores: Pearson correlation coefficient (PCC) [24], Spearman rank correlation (SRC) [28], mutual rank (MR) [25], and Euclidean distance (ED) [29]; and a supervised method: metric learning for co-expression (MLC) [21]. Each method is combined with the GBA strategy to predict GO terms, as described in Text S1 of supplementary information (SI). We evaluate the function prediction performance by four metrics: (1) Maximum F1-score (*Fmax*) [23], (2) area under the precision-recall curve (*AUPR*) [30], (3) weighted average *Fmax* (*WAFmax*), and (4) weighted average *AUPR* (*WAAUPR*) (see Equations 11-13 in “Materials and Methods”). **Figure 2 (A-B)** list the *WAFmax* and *WAAUPR* values on the test dataset of 5,754 genes from 8 species (human, mouse, arabidopsis, rat, fly, budding yeast, fission yeast, and nematoda, see “Materials and Methods”) for the six methods, where the p-values between TNP and the other five methods in Student’s t-test [31] for *WAFmax* and *WAAUPR* are summarized in Table S1 of SI. In addition, the performances of the six methods for each individual species are summarized in Figures S1-S2 and Table S2 and discussed in Text S2 of SI.

**Figure 2.**
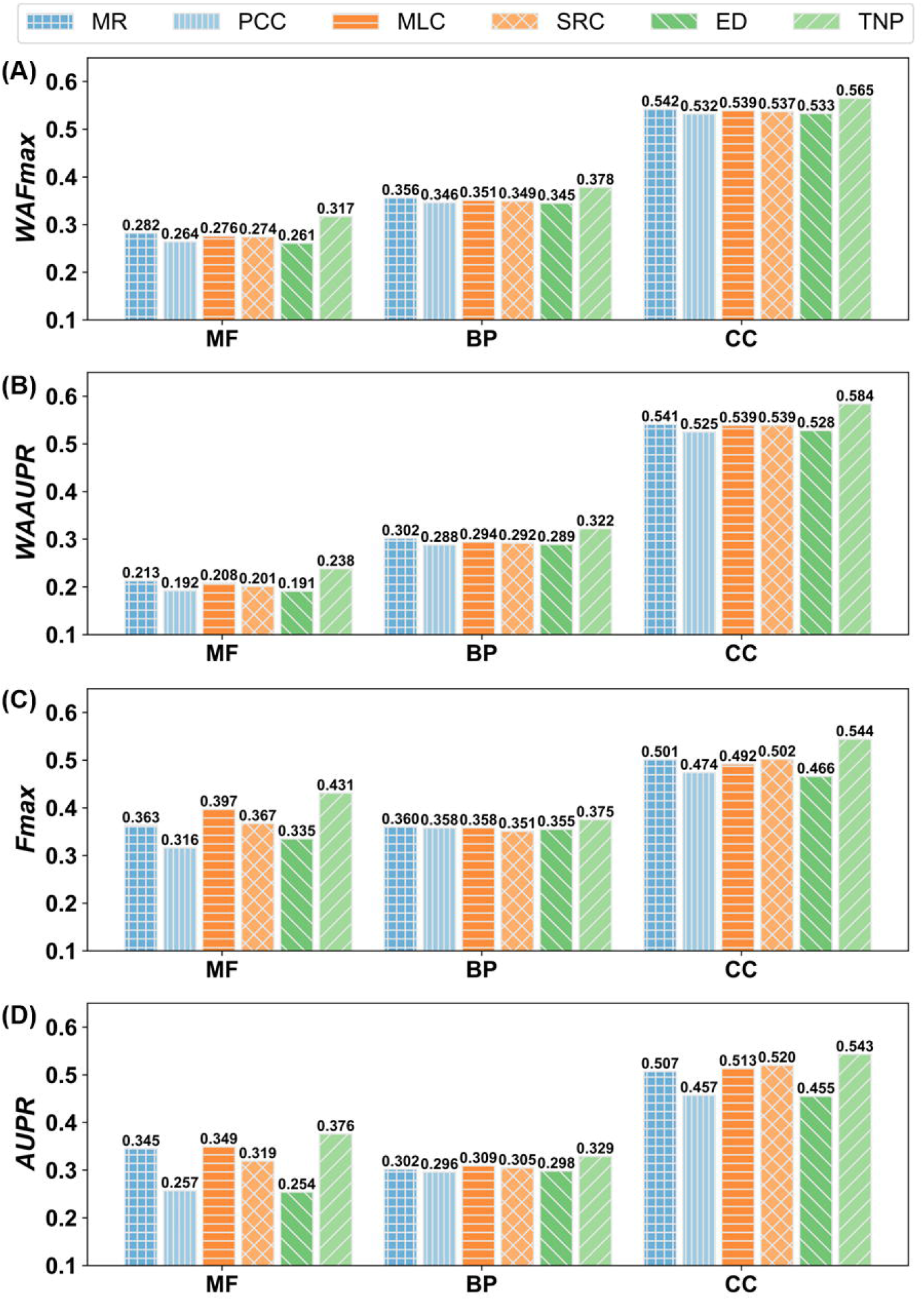
Comparison of six transcript expression profile-based GO prediction methods (key at top) **A-B**. The *WAFmax* and *WAAUPR* values on the test datasets of 8 species. **C-D**. The *Fmax* and *AUPR* values on 98 non-coding test genes. MR: mutual rank. PCC: Pearson correlation coefficient. MLC: metric learning for co-expression. SRC: Spearman rank correlation. ED: Euclidean distance. TNP (triplet network-based pipeline).

From **Figure 2 (A-B)** and Table S1, it can be observed that the accuracy of TNP is significantly higher than the other five methods in deducing function from gene expression data. Specifically, the improvements of *WAFmax* values between TNP and the second-best performer, MR, are 12.4% (= (0.317-0.282)/0.317×100%), 6.2%, and 4.2% for MF, BP, and CC, respectively, all with p-value significantly below 0.05. At the same time, TNP achieves an increase of *WAAUPR* values by 11.7%, 6.6%, and 7.9%, respectively, compared to MR for the three GO aspects. Moreover, TNP and ED separately show the best and worst performances among six methods, although they use the similar metric functions (TNP uses the square of Euclidean distance). These data suggest that the GO recognition accuracy can be improved via feature space mapping when coupled with triplet network learning. In addition, our result shows that MR obtains higher values of *WAFmax* and *WAAUPR* than PCC for each GO aspect; this is consistent with the fact that MR has replaced PCC as the new co-expression measure in current co-expression databases [32].

In **Figure 2 (C-D)**, we further compare the performances of the six expression profile-based methods on a subset of 98 non-coding genes in the test datasets of the 8 species, where TNP outperforms again all other five methods. Specifically, based on *Fmax*, TNP achieves 8.6%, 4.2%, and 8.4% improvements compared to the second-best performer for MF, BP, and CC, respectively. At the same time, the corresponding increases of *AUPR* values are 7.7%, 6.5%, and 4.4%, respectively, for the three GO aspects.

In addition, we used the human data to examine the influence of the different characteristics of gene-expression data on the prediction performance of TNP. First, as illustrated in Table S3 and Figure S3 and discussed in Text S3 of SI, the number of expression samples (i.e., the dimension of expression profile vector) is not critical to the TNP performance. In fact, the *Fmax* values of TNP trained on 30% of the expression samples are only slightly (i.e., by 1.3%, 1.5% and 0.7%) lower than those trained on all the data for MF, BP and CC, respectively. This is probably due to the inherent redundancy among different samples for the same species. To illustrate this point, we further compared TNP with and without its principal component analysis (PCA) [33] procedure, which was used to reduce redundant information of expression samples (see “Materials and Methods”). The result showed that skipping PCA leads to a clear and consistent drop of the performance (Figure S3), which confirms the negative impact of data redundancy on the performance. Finally, we found that there is no strong correlation between the function prediction accuracy of each gene (in terms of gene-level F1-score) and its mean expression level, with a neglectable PCC value ranging from -0.026 to 0.173 for all GO aspects (Figure S4 of SI).

### Expression similarity has closer correlation with functional similarity in embedding feature space than original feature space

One important component of TNP is feature space mapping, in which the expression score calculations are transferred from the original feature space to the embedding feature space (**Figure 1 (A-B)**). To examine the impact of the feature space mapping on GO prediction, we deigned and executed the following test.

For a query gene, we first rank all genes in a training dataset in descending order of the expression similarity between training gene and query, and select the top *K* (*K* = 100) genes as templates. In the original feature space, the expression similarity is measured by MR, PCC, MLC, SRC, and ED, respectively. In the embedding space, the expression similarity for TNP is calculated by the square of the Euclidean distance. Then, the weighted functional similarity (*WFS*), between templates and query, can be calculated as:

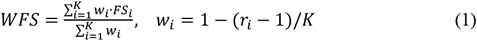

where *w*_*i*_ and *r*_*i*_ are the weight and rank, respectively, for *i*-th template, and *FS*_*i*_ is the functional similarity between the *i*-th template and query measured by F1-score between their experimental GO terms (see Text S4 of SI). Finally, the average weighted functional similarity (*AVG*_*WFS*) for all test genes is calculated by

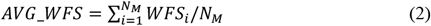

where *N*_*M*_ is the total number of test genes in the *M* species used here. A higher value of *AVG*_*WFS* indicates a closer correlation between expression similarity and functional similarity.

**Figure 3** shows the *AVG*_*WFS* values of six measures for three GO aspects in the 8 species. For each GO aspect, we find that TNP achieves the highest *AVG*_*WFS* among six measures. More specifically, the *AVG*_*WFS* values of TNP are 27.4%, 11.1%, and 7.9% higher than those of the second-best performer, MR, for MF, BP, and CC, respectively. Moreover, the *AVG*_*WFS* values of six measures in each individual species are listed in Figure S5 of SI, where TNP outperforms again other measures in all GO aspects.

**Figure 3.**
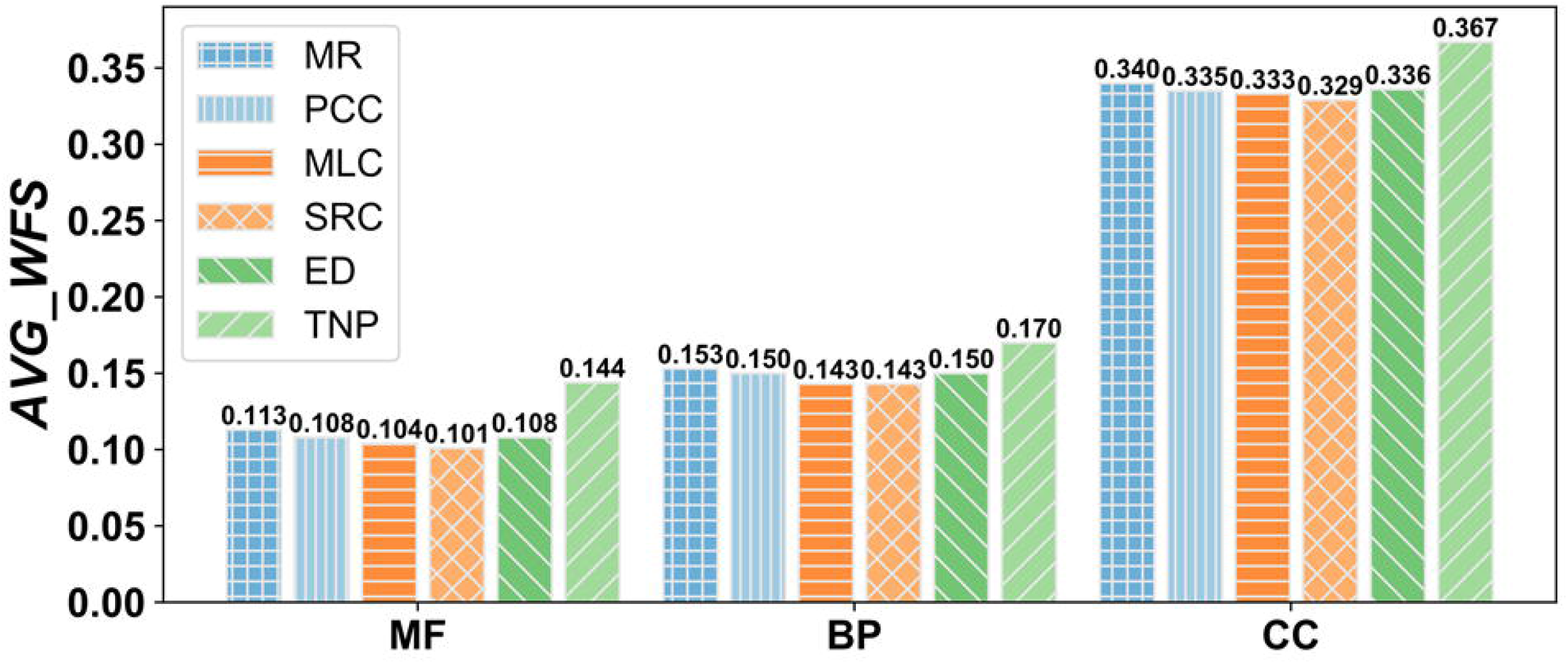
Comparison of the *AVG*_*WFS* values by six transcript expression profile-based GO prediction methods on 8 test species.

As an illustrative, we list in Figure S6 a scattering plot of F1-score versus weight for a non-coding gene MIRLET7C (Entrez ID: 406885) in the test dataset of human species. Here, we use three measures, TNP, MR and PCC, to select 100 templates with the highest expression similarity to the query. The expression similarity for different measures can be normalized as weight (*w*_*i*_ in Equation 1). The functional similarity is assessed by F1-score of experimental GO terms between two genes. It can be seen that TNP achieves a higher weighted functional similarity (*WFS*) value than both MR and PCC for each GO aspect, because it selects more templates which have a higher expression similarity (or weight) and functional similarity (F1-score) with the query than the two control measures. Since the data from TNP are directly taken from the embedding space after triplet-network training, these results suggest that the expression similarity for TNP in the embedding feature space has closer correlation with functional similarity compared to the other measures in the original space.

### Protein homology inference and triplet-network based expression make most important contributions on TripletGO prediction

To examine the contributions of four component methods in TripletGO, we compare the performances of four individual methods, including expression profile-based GO prediction (EPGP) by TNP, genetic sequence alignment-based GO prediction (GSAGP), protein sequence alignment-based GO prediction (PSAGP), and naïve-based GO prediction (NGP), and five combination methods, including GSAGP+PSAGP+NGP (GPN), EPGP+PSAGP+NGP (EPN), EPGP+GSAGP+NGP (EGN), EPGP+GSAGP+PSAGP (EGP), and EPGP+GSAGP+PSAGP+NGP (EGPN=TripletGO) (see “Materials and Methods”). To be fair, we optimized the confidence scores of the combination methods using the same network in **Figure 1C. Figure 4 (A-B)** list the *WAFmax* and *WAAUPR* values of all nine methods on the test datasets of 8 species, where the p-values between EGPN and the other eight methods in Student’s t-test for *WAFmax* and *WAAUPR* are listed in Table S4 of SI. In addition, the performances of all nine methods for each individual species are summarized in Figures S7-S9 and Table S5 and discussed in Text S5 of SI.

**Figure 4.**
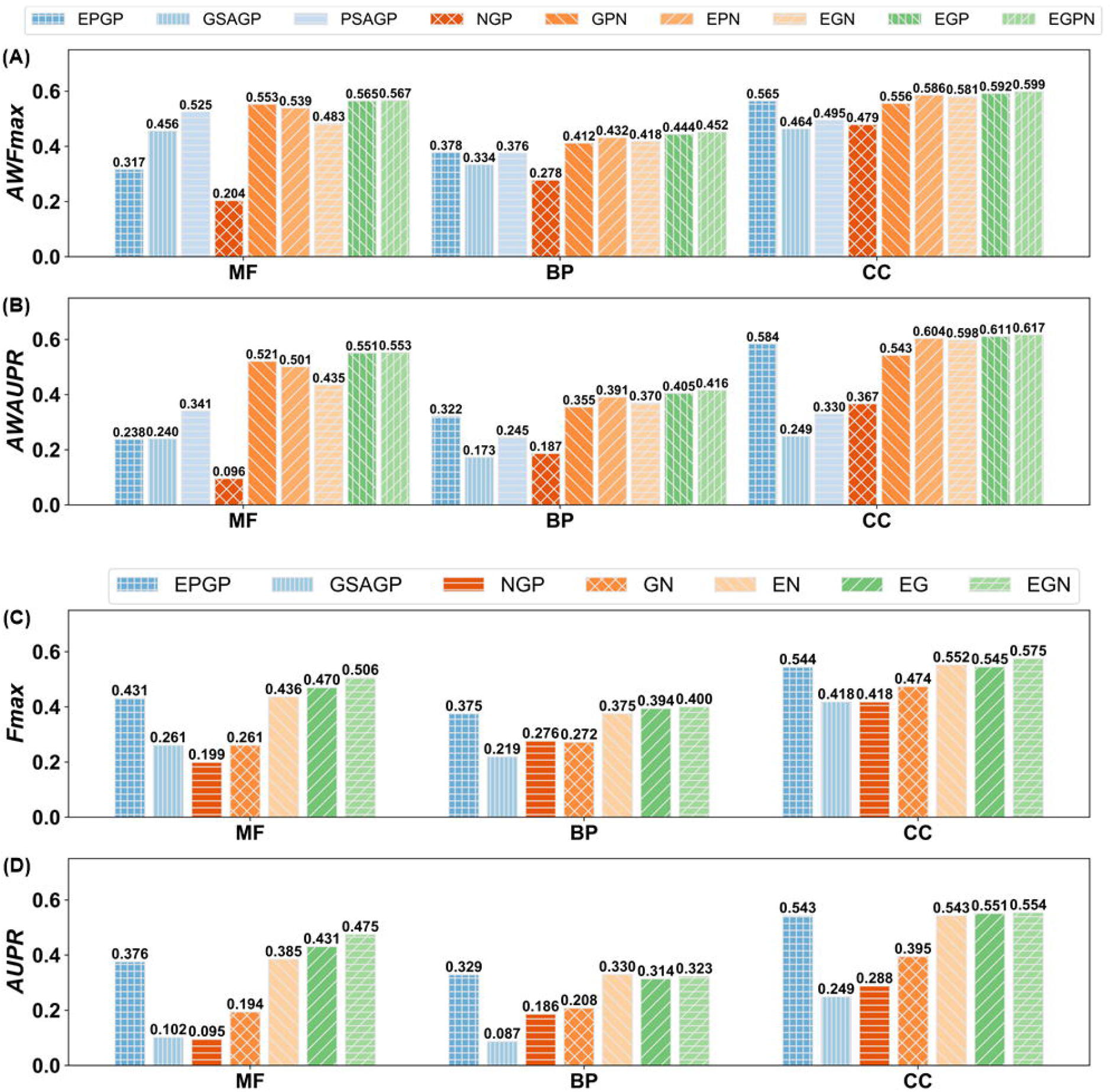
Comparison of GO prediction results using different methods. **A-B**. The *WAFmax* and *WAAUPR* values on the test datasets of 8 species for nine GO prediction methods. **C-D**. The *Fmax* and *AUPR* values on 98 non-coding genes for seven GO prediction methods.

From the data in **Figure 4 (A-B)** and Table S4, we can conclude that each individual method contributes to improving the TripletGO prediction performance. Specifically, the *WAFmax* and *WAAUPR* values of EGPN are much higher than the corresponding values by each of four individual methods. Importantly, the performance of EGPN is also significantly better than that of the other four combination methods. In terms of *WAFmax*, for example, EGPN gains 6.7%, 4.0%, 9.5%, and 1.1% average improvements for three GO aspects in comparison with GPN, EPN, EGN, and EPG, respectively. At the same time, the corresponding average increases of *WAAUPR* are 12.3%, 6.3%, 14.2%, and 1.4%, respectively. The first and second largest increases are caused by adding PSAGP to EGN and adding EPGP to GPN, respectively. In addition, among the four individual methods, PSAGP and EPGP achieve the best performance for MF, BP, and CC, respectively. These data demonstrate the importance of the protein-level homology inference and triplet-network based expression, respectively, to the TripletGO prediction.

We further investigate the contributions of proposed methods for non-coding genes. Since non-coding genes have no available prediction results in PSAGP, we compare the performances of three individual gene-level methods and four combination methods, including GSAGP+NPG (GN), EPGP+NPG (EN), EPGP+GSAGP (EG), and EPGP+GSAGP+NGP (EGN=TripletGO). **Figure 4 (C-D)** illustrate the *Fmax* and *AUPR* values of seven GO prediction methods for 98 non-coding genes. The p-values between EGN and other six methods in student’s t-test for *Fmax* and *AUPR* are shown in Table S6 of SI. Again, we can see that each of three gene-level methods helps to improve accuracy of GO prediction for non-coding genes. On the basis of *Fmax*, for example, EGN achieves the best performance among seven methods. Specifically, EGN gains 7.7%, 1.5%, and 4.2% improvements for MF, BP, and CC, respectively, compared to the second-best performer. With respect to *AUPR*, although EGN shows a slightly lower value than EPGP and EN in BP, it achieves the best performance for MF and CC. In addition, EPGP significantly outperforms other two individual methods through all score metrices, which highlights again the importance of the transcript expression component to the TripletGO prediction.

### Comparison of TripletGO with existing gene function prediction methods

We compare TripletGO with two most recently developed gene function prediction approaches, i.e., GENETICA [7] and GeneNetwork [34], which are both based on expression profiles. Different from our work, these two approaches are designed at the term-centric level. Specifically, for a GO term *Q*_*i*_, each gene is labeled as “1” or “0”, where “1” means this gene is associated with *Q*_*i*_ in the experimental annotation. Then, each gene is assigned with a confidence score for *Q*_*i*_ using leave-one-out strategy. Finally, the area under receiver operating characteristic curve (AUROC) is used to evaluate the prediction performance of *Q*_*i*_ by combining the confidence scores and labels for all genes. In light of this, our models are compared with GENETICA and GeneNetwork by term-centric evaluation.

Between our work and GENETICA, there are 287 MF, 1340 BP, and 186 CC terms in common for human. As for mouse, there are 149, 1230, and 128 common terms for MF, BP and CC, respectively (see Text S6 of SI). **Figure 5 (A-B)** plot the distributions of AUROC values by GENETICA, TNP, and TripletGO on three GO aspects in human and mouse, respectively. Figure S10 (A-B) in SI show the mean and median AUROC values for three methods. While TNP and GENETICA are both expression profile-based models, the former shows a better performance than the latter. In human, TNP achieves 16.5%, 10.8%, 14.4% increases of the mean AUROC for MF, BP, and CC, respectively, compared to GENETICA. At the same time, the corresponding increases of median AUROC are 23.0%, 8.5%, and 12.7%, respectively. As for mouse, although TNP gains a slightly lower median AUROC than GENETICA in CC, it achieves significant improvements of the corresponding measures in MF and BP. In addition, that TripletGO shows a significantly better performance than both TNP and GENETICA, mainly because it integrates complementary information from sources other than expression profiles.

**Figure 5.**
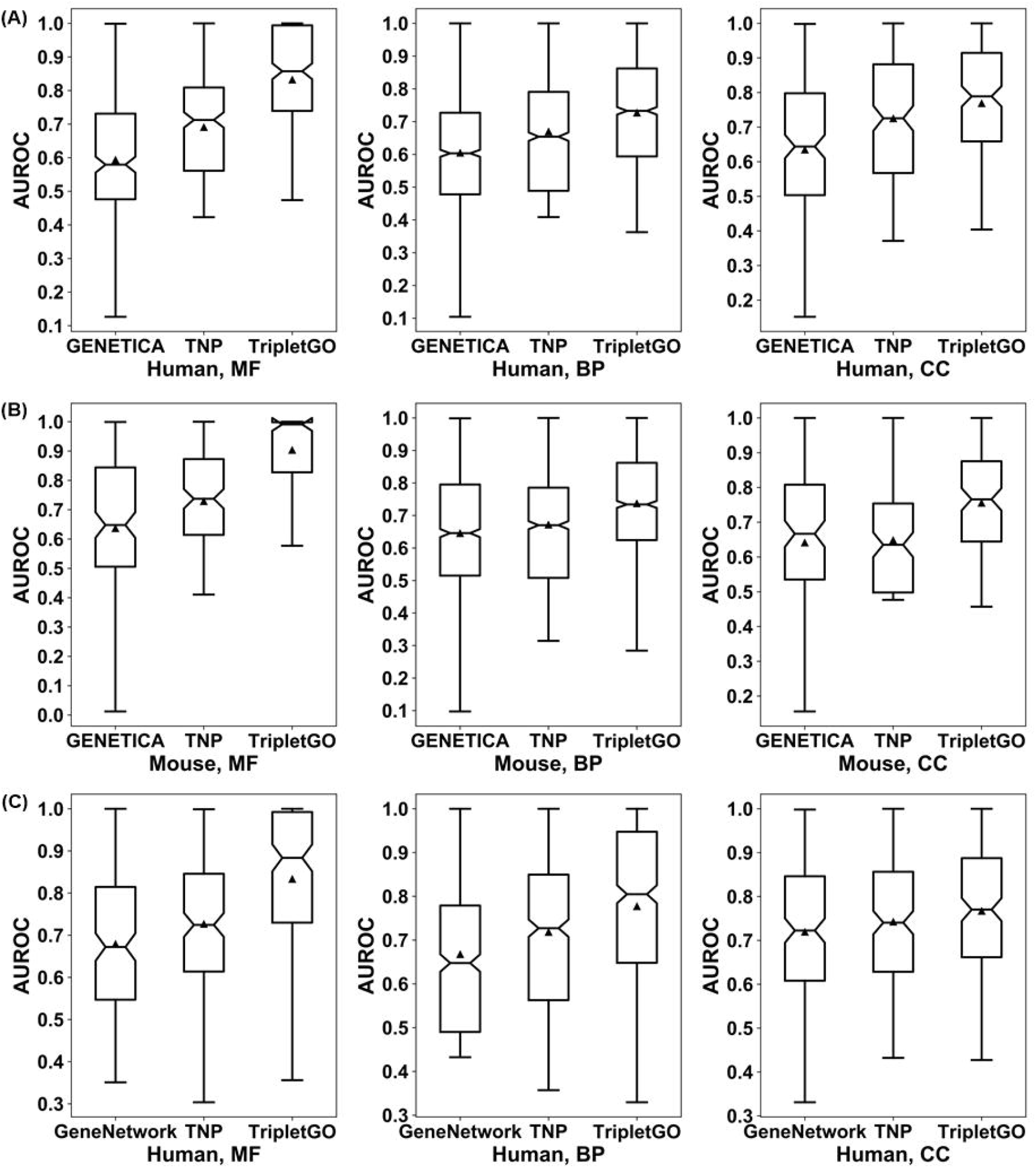
Comparison of AUROC values of three GO aspects by different methods on the common dataset. **A**. GENETICA, TNP and TripletGO on human. **B**. GENETICA, TNP and TripletGO on mouse. **C**. GeneNetwork, TNP and TripletGO on human. In each box, the median line and triangle represent the median and mean AUROC values, respectively.

There are 165, 522, and 182 common terms for MF, BP, and CC, respectively, in human genes between our work and GeneNetwork (see Text S7 of SI). **Figure 5C** shows the AUROC distributions of three GO aspects for GeneNetwork, TNP, and TripletGO, respectively, where Figure S10C in SI illustrates the mean and median AUROC values of three models. For each GO aspect, TNP shows higher mean and median AUROC values in comparison with GeneNetwork, while TripletGO outperforms both due to the integration of additional information from sequence homology alignments and prior statistics of Gene-GOA databases.

In Text S8 and Figure S11 of SI, we made a further comparison of our methods with GENETICA and GeneNetwork in the gene-center level based on *Fmax* and *AUPR*, where a similar trend (i.e., TripletGO and TNP outperform the control methods) but with more significant distinctions between the methods can be seen as the term-centric comparisons.

### Testing on CAFA3 targets

We further tested our methods on the dataset of the third CAFA challenge (CAFA3). The entire CAFA3 dataset consists of 66,841 training and 3,328 test proteins [23] from 23 species. Since some targets have no available gene expression data, we only benchmarked our methods on the 2,433 CAFA3 test proteins whose coding genes are originated from 7 species (human, mouse, arabidopsis, rat, fly, budding yeast and fission yeast) and have available expression profiles in COXPRESdb [32] or ATTED-II [25] databases. It should be noted that we did not find any test proteins with expression data from nematoda species. The details of training and test datasets for the 7 species are summarized in Table S7 of SI. For each species, we randomly select 90% training samples to re-train the TNP model and the remaining training samples are used to optimize the parameters of the model. Moreover, for GSAGP, PSAGP, and NPG, the entire CAFA3 training dataset is used to construct the corresponding template databases and prior probabilities of GO terms.

**Figure 6 (A-B)** summarizes the performance of six transcript expression profile-based GO prediction methods on the 2,433 test proteins, where the p-values between TNP and the other five methods in Student’s t-test [31] for *Fmax* and *AUPR* are listed in Table S8 of SI. It can be found that TNP shows better performance than other five methods for all GO aspects. Compared to the second-best performer (MR), TNP achieves 10.7% and 11.2% average improvements on three GO aspects for *Fmax* and *AUPR*, respectively, where the *p*-value is statistically significant for all the comparisons except for *AUPR* on MF (1.41E-01) and CC (8.01E-01). The performances of the six methods for each individual species are summarized in Figure S12 and Table S9, where TNP achieves the highest values of *Fmax* and *AUPR* among six methods for each GO aspect in most species (see discussion in Text S9 of SI).

**Figure 6.**
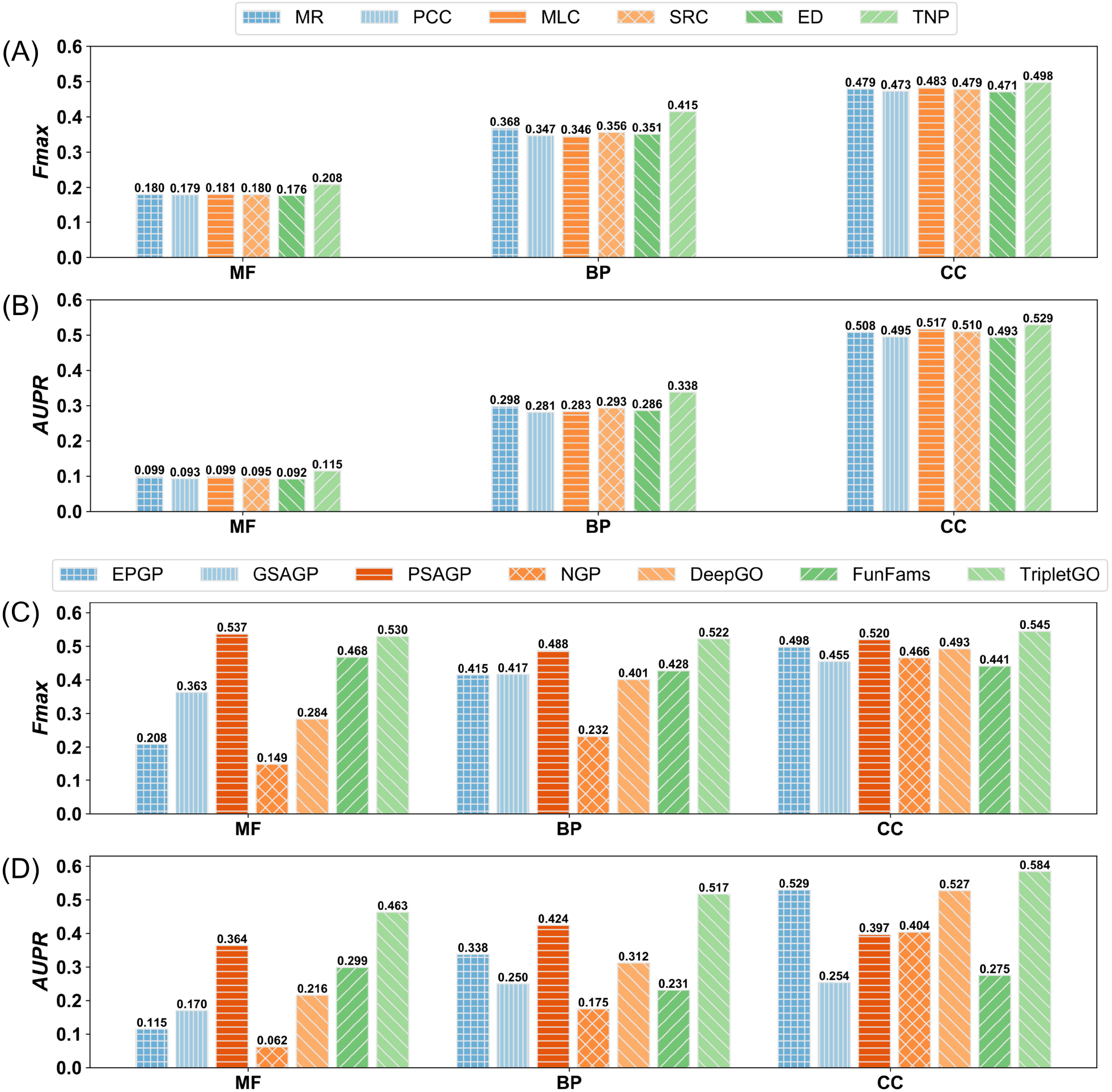
Performance comparison on 2,433 test proteins of 7 species from CAFA3 benchmark dataset. **A-B**. The *Fmax* and *AUPR* values for six transcript expression profile-based GO predictions. **C-D**. The *Fmax* and *AUPR* values for five proposed GO prediction methods and two existing GO prediction methods.

We further benchmarked five proposed GO prediction methods, including four individual methods (i.e., EPGP, GSAGP, PSAGP and NGP) and their combination (i.e., TripletGO), on the 2,433 CAFA3 test proteins. In addition, we included two third-party protein function prediction methods (DeepGO [10], FunFams [11]) which are the only methods that we found to have downloadable programs and allow us to test independently on our selected CAFA dataset. Meanwhile, they represent two typical types of protocols: while DeepGO is a machine-learning based method combining convolutional neural network with protein sequence encoding, FunFams is a template-based method and searches the functional templates using protein family information. Among them, FunFams is one of the top-performing methods and ranked at 2/4/9 position in MF/BP/CC aspects with respect to *Fmax* in the CAFA3 experiment [23]. To make a fair comparison between template-based and non-template-based methods, we used the pre-set cutoff (i.e., *t*_1_ = 60 % and *t*_2_ = 30 %, see “Materials and Methods”) to exclude close homologies when running GSAGP and PSAGP; however, we did not exclude any homologies for the third-party programs and ran them under the default setting.

**Figure 6 (C-D)** summarizes the performance of seven GO prediction methods, where the p-values between TripletGO and the other six methods in Student’s t-test [31] are listed in Table S10 of SI. We found that the composite GO prediction method, i.e., TripletGO, achieves a significantly better performance than other six GO prediction methods in all GO aspects, including both the third-party (FunFams and DeepGO) and the component methods of TripletGO, demonstrating again the advantage of integrating gene expression and sequence profile-based approach to function predictions.

### Case studies

As illustrations, we selected two genes from the human genome: GALNT4 (protein-coding gene, Entrez ID: 8693) and MIRLET7C (non-coding gene, Entrez ID: 406885), to examine the effects of different GO prediction methods. Here, each gene is associated with 12 GO terms for CC aspect from experimental annotations. **Table 1** summarizes the numbers of correctly predicted GO terms (i.e., true positives) and mistakenly predicted terms (i.e., false positives) in CC aspect for the two genes by ten different methods, including six individual gene-expression methods (MR, PCC, MLC, SRC, ED, and TNP), a gene sequence alignment method (GSAGP), a protein sequence alignment method (PSAGP), a naïve-based approach (NGP), and a composite approach (TripletGO). **Figure 7** plots the directed acyclic graph of GO terms in native annotation and the correctly predicted GO terms of ten methods for the two genes. Moreover, the incorrectly predicted GO terms (i.e., false positives) of each method are listed in **Table 2**. It should be noted that the predicted GO terms of each method are determined by its own cut-off value to achieve the highest *Fmax* value.

**Table 1.** The modeling results of ten GO prediction methods on two illustrative genes. TP and FP refer to true positive and false positive rates, respectively. Best performers are highlighted in bold fonts in each category.

| Gene | Measure | MR | PCC | MLC | SRC | ED | TNP | GSAGP | PSAGP | NPG | TripletGO |
| --- | --- | --- | --- | --- | --- | --- | --- | --- | --- | --- | --- |
| GALNT4 | TP | 8 | 8 | 7 | 7 | 8 | 8 | 7 | 7 | 7 | **10** |
|  | FP | 3 | 1 | 2 | 3 | 1 | **0** | 3 | 4 | 4 | **0** |
| MIRLET7C | TP | 1 | 1 | 3 | 6 | 1 | 7 | **8** | 0 | 6 | **8** |
|  | FP | 2 | 2 | 2 | 2 | 2 | 1 | **0** | **0** | 2 | **0** |

**Table 2.** The incorrectly predicted GO terms for ten GO prediction methods on two illustrative genes.

|  | GALNT4 | MIRLET7C |
| --- | --- | --- |
|  | False positives | False positives |
| MR | GO:0005654 GO:0005829 GO:0032991 | GO:0016020 GO:0005886 |
| PCC | GO:0005829 | GO:0016020 GO:0005886 |
| MLC | GO:0005829 GO:0032991 | GO:0016020 GO:0005886 |
| SRC | GO:0005654 GO:0005829 GO:0005886 | GO:0016020 GO:0005886 |
| ED | GO:0005829 | GO:0016020 GO:0005886 |
| TNP |  | GO:0005654 |
| GSAGP | GO:0005654 GO:0005829 GO:0005886 |  |
| PSAGP | GO:0031410 GO:0030133 GO:0097708 GO:0031982 |  |
| NGP | GO:0005829 GO:0032991 GO:0005634 GO:0005886 | GO:0016020 GO:0032991 |
| TripletGO |  |  |

**Figure 7.**
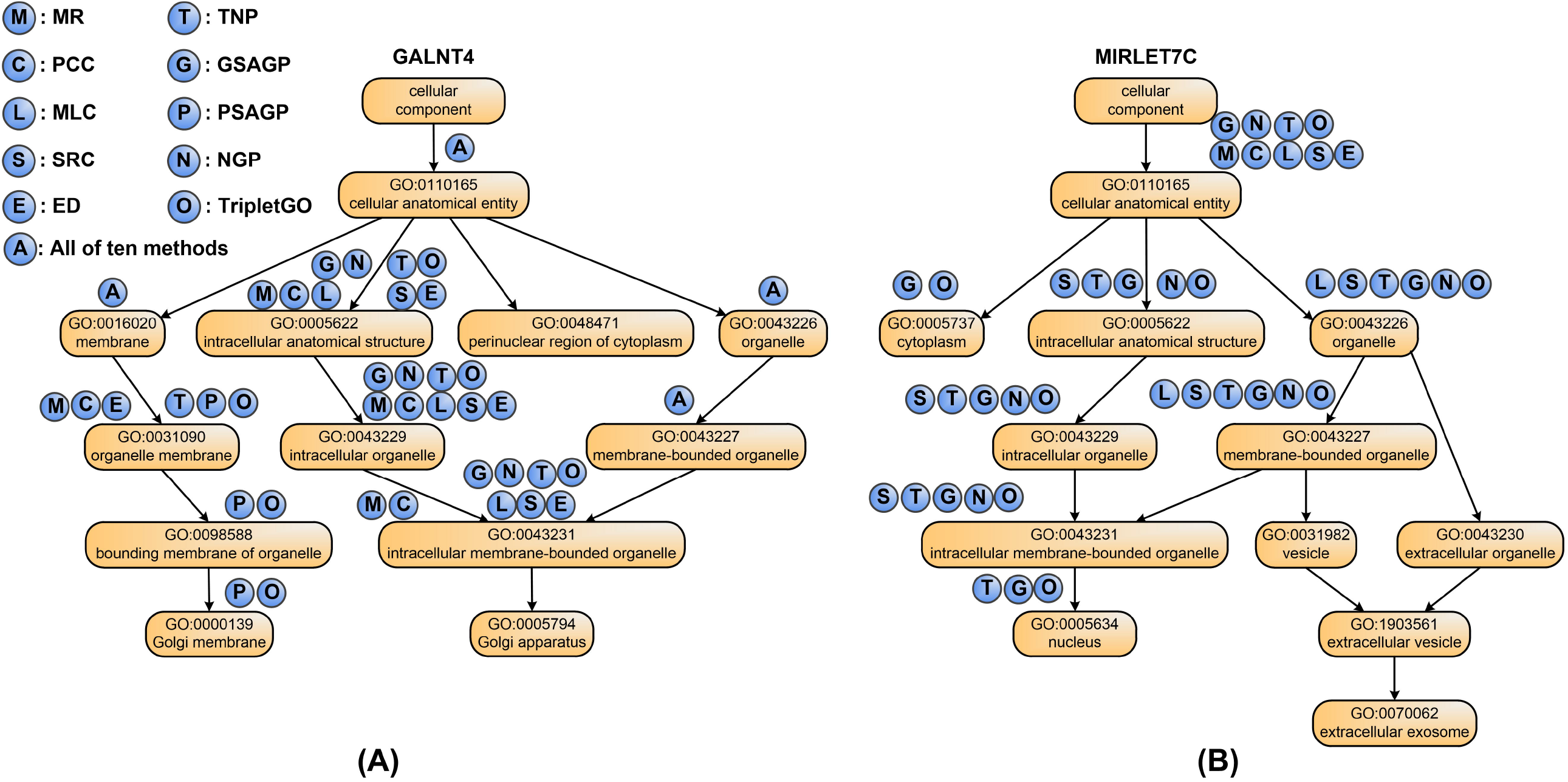
The directed acyclic graphs of 12 GO terms in the experimental annotation on two illustrative genes. **A**.GALNT4; **B**. MIRLET7C. The circles above each GO term refer to the prediction methods, where a circle filled with “X” on GO term “Y” indicates that method “X” can correctly predict term “Y”.

Several interesting observations can be made from the data. First, among six gene-expression based methods, TNP can correctly recognize the most GO terms with the least false positives for each gene. Moreover, all true positives for other five methods can be effectively identified by TNP. More importantly, for gene MIRLET7C, TNP correctly recognizes 1 additional GO term, i.e., GO:0005634, which are missed by the other five methods. This observation shows that TNP can predict gene function in a more precise level, because it successfully identifies some children GO terms, in which other expression-based methods fail.

Second, the combination of complementary methods increases both coverage and accuracy of the TripletGO models. In gene GALNT4 (see **Figure 7A**), the four component methods (TNP, GSAGP, PSAGP, and NGP) hit 10 true positives in total, which is more than that by each individual method, indicating that the component methods derive complementary information from different sources. By taking the combination, TripletGO achieves the highest coverage with TP=10 and the lowest false positive rate (FP=0). Sometimes, one component method can cover all true positives predicted by other methods. For example, for gene MIRLET7C (see **Figure 7B**), all true positives of TNP and NGP are covered by GSAGP. Even in this case, the final TripletGO accuracy is not degraded by the inclusion of a less accurate method, where TripletGO shares the same performance with the best individual method by GSAGP.

In addition, to further explain what is considered as a positive prediction regarding the hierarchy, we choose GALNT4 as an illustrative example and list the confidence scores of all the candidate GO terms by TripletGO in Text S10 of SI, where the 10 GO terms whose confidence scores are higher than the cut-off value (0.35) have been precited as positives. In Figure S13, we plot the directed acyclic graph of the 10 predicted GO terms with corresponding confidence scores. It can be found the confidence scores of the parent terms are higher than the scores of their children, following the post-processing Equation 10 in “Materials and Methods”.

## Conclusions

We developed a new method, TripletGO, to predict the function of both protein-coding and non-coding genes by the integration of four gene-expression and protein homology inference pipelines. The large-scale benchmark tests on 5,754 non-redundant genes from a set of 8 species demonstrated that TripletGO consistently achieved significant improvements in comparison with other state-of-the-art gene function prediction methods. Detailed analysis showed that the major advantage of TripletGO stems from two aspects. First, the new triplet network-based algorithm, when coupled with feature space mapping, efficiently recognizes functional patterns from transcript expression profiles. Second, the combination of multiple complementary pipelines, especially those with protein-level homology inference and transcript expression profile, significantly improves the coverage and accuracy of the gene function annotations.

Despite the encouraging performance, there is still considerable room for further improvements. First, the TNP needs large amounts of gene expression data with GO annotation to train the prediction model. For some species, such as dog and chicken as listed in Table S11 of SI, the number of genes with GO annotation is very limited, and for many other species of interest no such data are available. As a result, we cannot train prediction models using the TNP from expression profiles for these species. Therefore, an extended TNP model by normalizing expression profiles across different species may help solve the issue, as well as further improve the overall accuracy of the current approach. Second, the confidence scores of the four individual methods are integrated as a consensus score by a simple one-layer neural network, where an advanced machine-learning approach may help better integrate confidence scores. Meanwhile, new GO prediction methods considering other biological aspects, such as protein-protein interactions and protein/nucleic acid structures, will help improve both the coverage and accuracy of the current gene function annotation algorithms. Studies along these lines are under progress.

## Materials and methods

### Datasets

We collected all 78,170 genes with GO annotation via experimental determination from National Center for Biotechnology Information (NCBI) [35] to construct the Gene-GOA database (Text S11 of SI). The genes in Gene-GOA were used for both template database construction and for prior probability construction in the naïve statistics-based GO modeling.

To evaluate the proposed methods, we collected 57,584 genes from 8 species by the following procedures: (1) we downloaded all of 300,977 genes with expression profiles determined by microarray [36] for 20 species from COXPRESdb [32] and ATTED-II [25] databases. For each species, the total number of genes with functional annotation in Gene-GOA is shown in Table S11 of SI. Then, we select the 8 species with the most genes with GO annotation among the 20 species, to construct benchmark datasets. (2) For each species, we randomly select 85% of genes with GO annotation as the training dataset, and 5% genes as the validation dataset, which are separately used to construct machine learning-based models and optimize the parameters of models. The remaining 10% genes are used as the test dataset to assess the performance of models. As a result, there are 48,954, 2,876, and 5,754 genes in training, validation, and test datasets, respectively, for the 8 species in total, as summarized in Table S12 of SI.

### Expression profile-based GO prediction by TNP

In expression profile-based GO prediction (EPGP), a triplet network [27] is used to measure the similarity of expression profiles, as shown in **Figure 1A**. The input is a triplet variable (*anc, pos, neg*), where *anc* is an anchor (baseline) gene, *pos* is a positive gene with the same function of *anc*, and *neg* is a negative gene with the different function of *anc*. First, the expression profile of each gene is mapped from the original feature space to an embedding space using the same deep neural network. Next, the expression dissimilarity between two genes in embedding space is measured by Euclidean distance [29] of the mapped expression profiles. Finally, the triplet loss function is designed to associate expression similarity with functional similarity:

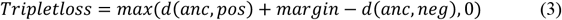

where *d*(*anc, pos*) is the Euclidean distance between anchor and positive genes in embedding space, *d*(*anc, neg*) is the distance between anchor and negative genes, and *margin* is a pre-set positive value. Here, the minimization of the triplet loss requests for the maximization of *d*(*anc, neg*) − *d*(*anc, pos*). In the ideal case, *tripet loss* = 0 when *d*(*anc, neg*) ≥ *d*(*anc, pos*) + *margin*, which indicates substantially higher similarity (lower distance) of the anchor genes to the positive genes than to the negative genes.

It has been demonstrated that cross-entropy loss [37] helps to improve the performance of triplet network [38, 39]. Therefore, we further combine the triplet loss with the cross-entropy loss in the TNP to predict gene function from expression profiles. The overall workflow of TNP is depicted in **Figure 1B**, which contains two stages.

### Training stage of TNP

#### Procedure I: expression profile normalization

In a training dataset, the expression profiles of all *m* genes are represented as a matrix ***E*** = (*e*_*ij*_)_*m*×*l*_, where *l* is the number of experimental samples in microarray technology [36], and *e*_*ij*_ is the expression value of the *i*-th gene on the *j*-th sample. Each row of ***E*** can be viewed as the expression profile of a gene. To reduce noise and computing cost, the matrix ***E*** is transformed into a normalized matrix 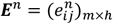 (*h* < *l*) by performing z-score normalization [40] and principal component analysis (PCA) [33].

#### Procedure II: expression profile mapping using a neural network

The normalized expression profiles are mapped from the original feature space to an embedding space using a neural network. Specifically, the normalized matrix ***E***^*n*^ is fed to a deep fully connected block (DFCB) with *N* layers to output an embedding matrix 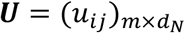, where *d*_*N*_ is number of neurons in the *N*-th layer. Then, L2-normaliztion is executed on ***U*** to obtain a normalized matrix 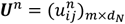, where 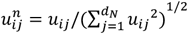. Each row of ***U***^*n*^ can be viewed as the expression profile of a gene in the embedding space.

At the same time, an output layer *L*_*O*_ with sigmoid activation function [41] is fully connected with DFCB to output a score matrix ***S*** = (*s*_*ij*_)_*m*×*r*_, where *r* is the number of GO terms in the training dataset, and *s*_*ij*_ is the confidence score that the *i* -th training gene is associated with the *j*-th GO term. Then, we calculate triplet loss and cross-entropy loss based on matrix ***U***^*n*^ and score matrix ***S***, respectively.

#### Procedure III: loss function calculation and network optimization

We use the “batch on hard” strategy [42, 43] to calculate the triplet loss:

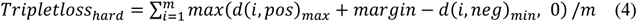

where *d*(*i, pos*)_*max*_ (or *d*(*i, neg*)_*min*_) is the maximum (or minimum) value of distances between the *i*-th gene and all positive (or negative) genes with same (or different) function of the *i*-th gene in embedding space. The distance between the two genes (*i, j*) is measured by 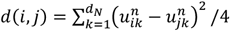, where the division factor of 4 is introduced to normalize *d*(*i, j*) into the range of [0,1], i.e., 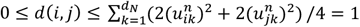. Moreover, two genes are considered to have the same function if their functional similarity is larger than a cut-off value *c*_*f*_. The functional similarity of two genes is measured by the F1-score between their GO terms, as shown in Text S4 of SI.

The cross-entropy loss is calculated as:

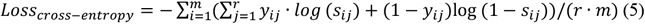

where *y*_*ij*_=1 if the *i*-th gene is associated with the *j*-th GO term in the experimental function annotation; otherwise, *y*_*ij*_=0.

The final training loss in TNP is the combination of triplet loss and cross-entropy loss:

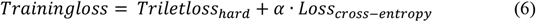

where *α* is a trade-off value. Finally, we minimize training loss to optimize neural network by Adam optimization algorithm [44].

### Prediction stage of TNP

The input is a query gene with expression profile vector ***e***^*q*^, and the output is a confidence score vector ***s***, including the confidence scores of *r* GO terms for query. First, z-score normalization and PCA are orderly executed on ***e***^*q*^ to obtain a normalized vector 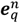, used as the input of DFCB. Then, we execute L2-normalization on the output of DFCB to obtain a normalized embedding vector ***u***^*q*^. Next, a distance rank-based strategy (see details in Text S12 of SI) is executed on the normalized embedding matrix of training genes (***U***^*n*^) and ***u***^*q*^ to generate a confidence score vector ***s***^*t*^. At the same time, the output layer *L*_*O*_ outputs another score vector ***s***^*c*^ by sigmoid function mapping. The final score vector ***s*** is the combination of two vectors:

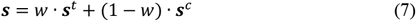

where *w* is a trade-off value and ranges from 0 to 1.

### Genetic sequence alignment-based GO prediction (GSAGP)

In GSAGP, we search the template genes, which have the similar sequences with query gene, from a genetic sequence database with GO annotation (GSD-GOA, see Text S13 of SI) for functional annotation.

For a query, we extract its RNA sequence from NCBI. Then, Blastn [45] is used to search the templates of query with e-value cutoff of 0.1 against GSD-GOA. To remove homology contamination, we exclude all homologous templates which have more than *t*_1_ sequence identity with the query. Finally, the remaining templates are used to annotate the query. Specifically, the confidence score that the query is associated with GO term *Q*_*i*_ is calculated as:

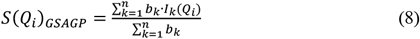

where *n* is the number of template genes, *b*_*k*_ is the bit-score of *k*-th template by Blastn; *I*_*k*_(*Q*_*i*_) = 1, if the *k*-th template is associated with *Q*_*i*_ in the experimental function annotation; otherwise, *I*_*k*_(*Q*_*i*_) = 0.

### Protein sequence alignment-based GO prediction (PSAGP)

In PSAGP, we select the template genes, whose coding proteins have similar sequence with that of the query, for GO functional annotation.

For a query gene, we map it as the corresponding coding protein sequence *P* in the UniProt database [46]. Then, Blastp [45] is used to search the template proteins of *P* with e-value cutoff of 0.1 against a protein sequence database (i.e., PSD, see Text S14 of SI), where homologous templates with a sequence identity above *t*_2_ to *P* are removed. Finally, the remaining templates are mapped back to genes in Gene-GOA to annotate query. The confidence score is calculated using the same scoring function as in GSAGP (i.e., Equation 8), where *b*_*k*_ is the bit-score of *k*-th template by Blastp.

### Naïve-based GO prediction (NGP)

In NGP, the confidence score that a query is associated with GO term *Q*_*i*_ can be calculated by the frequency of *Q*_*i*_ in Gene-GOA:

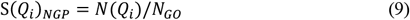

where *N*(*Q*_*i*_) is the number of genes associated with *Q*_*i*_, and *N*_*GO*_ is the number of genes with at least one annotation for the same GO aspect as *Q*_*i*_. This predictor can be thought of as a prior arising from the overall abundance of a particular annotation in Gene-GOA.

### Consensus TripletGO model

In TripletGO, the final GO model is a combination of the outputs of the above four pipelines as shown in **Figure 1C**. Here, the input is a genetic sequence with Entrez Gene ID, and the output is the confidence score for the predicted GO term. First, we extract the expression profile and coding protein sequence for query gene from COXPRESdb (or ATTED-II) database and UniProt database, respectively, using Entrez ID. Then, the expression profile, genetic sequence, and protein sequence are separately used as the inputs of EPGP, GSAGP, and PSAGP methods to output the confidence scores of GO terms. Moreover, NGP method is also used to calculate confidence scores. For a term *Q*_*j*_, its confidence scores by the four methods are serially combined as a vector used as the input of fully connected neural network to output the consensus score.

### Implementation and parameter settings of TripletGO

In EPGP, DFCB consists of two fully connected layers, each including 1,024 neurons with RELU activation function [47]. The remaining parameters of EPGP are listed in Table S13 of SI. In PSAGP, we use *t*_2_ =30% sequence identity as the cut-off to remove homologous protein templates, following previous studies [9]. To determine the homology cutoff for nucleotide sequences, we use three different machine learning models to fit the relationship between protein sequence identity and gene sequence identity, and it was found that a 30% protein sequence identity roughly corresponds to 60% genetic sequence identity, as shown in Text S15 and Figure S14 of SI. Therefore, we use *t*_1_ =60% sequence identity as cut-off to remove homologous templates in the GSAGP.

### Consideration of hierarchical relation for evaluation of GO annotation

The GO annotation is hierarchical [23]. Specifically, for both the ground truth and the prediction, if a protein (gene) is annotated with a GO term *Q*_*i*_, it should be annotated with the direct patent and all ancestors of *Q*_*i*_. To enforce this hierarchical relation, we follow CAFA’s rule and use a common post-processing procedure [10] for the confidence score of term *Q*_*i*_ in all GO prediction methods as follows:

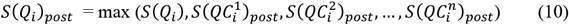

where *S*(*Q*_*i*_) and *S*(*Q*_*i*_)_*post*_ are the confidence scores of *Q*_*i*_ before and after post-processing, 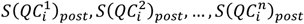 are the confidence scores of all direct children terms of *Q*_*i*_ after post-processing. This post-processing procedure enforces that the confidence score of *Q*_*i*_ is larger than or equal to the scores of all children.

### Evaluation metrices

Maximum F1-score (*Fmax*) and area under the precision-recall curve (*AUPR*) are used to evaluate the performance of proposed methods. *Fmax* is one of the most important evaluation metrics in CAFA [23, 48] and is defined as:

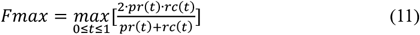

where *t* is a cut-off value of confidence score; *pr*(*t*) and *rc*(*t*) are precision and recall, respectively, with confidence score ≥ *t*:

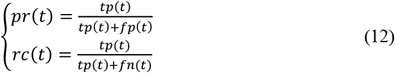

where *tp*(*t*) is the number of correctly predicted GO terms, *tp*(*t*) + *fp*(*t*) is the number of all predicted GO terms, and *tp*(*t*) + *fn*(*t*) is the number of GO terms in experimental function annotation. *AUPR* is a critical measure in multi-label prediction task [49] and ranges from 0 to 1.

The average performance of a method on multiple species is measured by weighted average *Fmax* (*WAFmax*) and weighted average *AUPR* (*WAAUPR*):

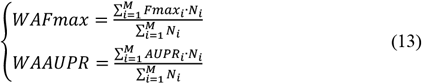

where *M* is the number of species, *Fmax*_*i*_ and *AUPR*_*i*_ are the *Fmax* and *AUPR* values, respectively, on the test dataset of the *i*-th species, and *N*_*i*_ is the number of test genes for the *i*-th species.

## Code availability

The online server, standalone package, and all benchmark datasets and libraries are available at https://zhanglab.ccmb.med.umich.edu/TripletGO/.

## CRediT author statement

Yang Zhang conceived and designed research; Yi-Heng Zhu wrote algorithm and performed experiment; Yi-Heng Zhu and Chengxin Zhang designed server; Chengxin Zhang and Yan Liu participated in discussion; Dong-Jun Yu and Yang Zhang supervised study; Yi-Heng Zhu and Yang Zhang wrote manuscript; Gilbert S. Omenn, Peter L. Freddolino and Dong-Jun Yu edited manuscript; All authors proofread and approved manuscript.

## Competing interests

The authors declare that they have no competing interests

## Acknowledgements

This work is supported in part by the National Natural Science Foundation of China (62072243 and 61772273 to Dong-Jun Yu), the Natural Science Foundation of Jiangsu (BK20201304 to Dong-Jun Yu), the Foundation of National Defense Key Laboratory of Science and Technology (JZX7Y202001SY000901 to Dong-Jun Yu), China Scholarship Council (201906840041 to Yi-Heng Zhu), the National Institute of Environmental Health Sciences (P30ES017885 to Gilbert S. Omenn), the National Cancer Institute (U24CA210967 to Gilbert S. Omenn), the National Institute of General Medical Sciences (GM136422, S10OD026825 to Yang Zhang), the National Institute of Allergy and Infectious Diseases (AI134678 to Peter L. Freddolino and Yang Zhang), and the National Science Foundation (IIS1901191, DBI2030790, MTM2025426 to Yang Zhang). This work used the Extreme Science and Engineering Discovery Environment (XSEDE), which is supported by National Science Foundation (ACI1548562). The work was done when Yi-Heng Zhu visited the University of Michigan.

## Supplementary material

### Supporting Figures

**Figure S1 The performance of six expression profile-based methods on the test datasets for 8 species**

**Figure S2 The precision-recall curves of six expression profile-based methods on the test datasets for 8 species**

**Figure S3 Variation curves of *Fmax* values of TNP and NON-PCA-TNP on the test dataset of human species versus the sampling ratios in expression data**

**Figure S4 The scattering plots of mean expression level versus F1-scores for 1470 human test genes by TNP**

**Figure S5 The *AVG***_***WFS* values of six measures for three GO aspects in 8 individual species**

**Figure S6 The scattering plots of weights versus F1-scores of 100 templates for the gene MIRLET7C over TNP, MR, and PCC**

**Figure S7 The *Fmax* values of nine GO prediction methods on the test datasets for 8 species**

**Figure S8 The *AUPR* values of nine GO prediction methods on the test datasets for 8 species**

**Figure S9 The precision-recall curves of five GO prediction methods on the test datasets for 8 species**

**Figure S10 Comparison of mean and median AUROC values of three GO aspects by different methods on the common dataset**

**Figure S11 Comparison of *Fmax* and *AUPR* values of three GO aspects by different methods on the common dataset**

**Figure S12 The performance of six expression profile-based methods for 7 species on CAFA3 test dataset**

**Figure S13 The directed acyclic graph of predicted GO terms with corresponding confidence scores for gene GALNT4 by TripletGO**

**Figure S14 The distribution of sequence identities for 10000 gene-gene pairs and 10000 mapped protein-protein pairs**

### Supporting tables

**Table S1 The p-values between TNP and other five expression profile-based methods for *WAFmax* and *WAAUPR***

**Table S2 The p-values between TNP and other five expression profile-based methods for *Fmax* and *AUPR* on 8 species**

**Table S3 The *Fmax* values of TNP and NON-PCA-TNP on the test dataset of human species for different sampling ratios in expression data**

**Table S4 The p-values between EGPN and other eight GO prediction methods for *WAFmax* and *WAAUPR* on the test datasets of 8 species**

**Table S5 The p-values between EGPN and other eight GO prediction methods for *Fmax* and *AUPR* on 8 species**

**Table S6 The p-values between EGN and other six GO prediction methods for *Fmax* and *AUPR* on 98 non-coding genes**

**Table S7 The details of training and test dataset for 7 species in CAFA3 dataset**

**Table S8 The p-values between TNP and other five expression profile-based methods for *Fmax* and *AUPR* on 2433 proteins of 7 species from CAFA3 test dataset**

**Table S9 The p-values between TNP and other five expression profile-based methods for *Fmax* and *AUPR* on CAFA3 test dataset for each of 7 species**

**Table S10 The p-values between TripletGO and other six GO prediction methods for *Fmax* and *AUPR* on 2433 proteins of 7 species from CAFA3 test dataset**

**Table S11 The numbers of genes with GO annotation of three aspects for 20 species**

**Table S12 The details of 8 benchmark datasets constructed in our work**

**Table S13 The values of *α, h, margin*, and *c***_***f***_ **on the benchmark datasets for 8 species**

### Supporting Texts

**Text S1 The procedures of GBA strategy for expression profile-based GO prediction**

**Text S2 The performances of six expression profile-based GO prediction methods for each individual species**

**Text S3 Exploring the influence of the characteristics of expression data on prediction performance for human species**

**Text S4 The functional similarity for genes**

**Text S5 The performances of nine GO prediction methods for each individual species**

**Text S6 Finding common genes and GO terms between our datasets and GENETICA’s datasets**

**Text S7 Finding common genes and GO terms between our datasets and GeneNetwork’s datasets**

**Text S8 Comparison with the existing gene function prediction models in gene-center level**

**Text S9 The performances of six expression profile-based GO prediction methods for each individual species on CAFA3 test dataset**

**Text S10 The confidence scores of the candidate GO terms for gene GALNT4 by TripletGO**

**Text S11 The construction procedures of Gene-GOA Text S12 Distance rank-based strategy**

**Text S13 The construction procedures of genetic sequence database with GO annotation**

**Text S14 The construction procedures of protein sequence database**

**Text S15 The relationship between protein sequence identity and genetic sequence identity**

## Notes

https://zhanglab.ccmb.med.umich.edu/TripletGO/

### Competing Interest Statement

The authors have declared no competing interest.

