## Supporting Information for "TripletGO: Integrating Transcript Expression Profiles with Protein Homology Inferences for High-Accuracy Gene Function Annotations"

#### Table of contents

##### Supporting Texts

**Text S1**. The procedures of GBA strategy for expression profile-based GO prediction.

**Text S2**. The performances of six expression profile-based GO prediction methods for each individual species.

**Text S3**. Exploring the influence of the characteristics of expression data on prediction performance for human species.

**Text S4**. The functional similarity for genes.

**Text S5**. The performances of nine GO prediction methods for each individual species.

**Text S6**. Finding common genes and GO terms between our datasets and GENETICA's datasets.

**Text S7**. Finding common genes and GO terms between our datasets and GeneNetwork's datasets.

**Text S8**. Comparison with the existing gene function prediction models in gene-center level.

**Text S9**. The performances of six expression profile-based GO prediction methods for each individual species on CAFA3 test dataset

**Text S10**. The confidence scores of the candidate GO terms for gene GALNT4 by TripletGO

**Text S11**. The construction procedures of Gene-GOA.

**Text S12**. Distance rank-based strategy.

**Text S13**. The construction procedures of genetic sequence database with GO annotation.

**Text S14**. The construction procedures of protein sequence database.

**Text S15**. The relationship between protein sequence identity and genetic sequence identity.

##### Supporting Tables

**Table S1**. The p-values between TNP and other five expression profile-based methods for $WAFmax$ and $WAAUPR$.

**Table S2**. The p-values between TNP and other five expression profile-based methods for $Fmax$ and $AUPR$ on 8 species.

**Table S3**. The $Fmax$ values of TNP and NON-PCA-TNP on the test dataset of human species for different sampling ratios in expression data.

**Table S4**. The p-values between EGPN and other eight GO prediction methods for $WAFmax$ and $WAAUPR$ on the test datasets of 8 species.

**Table S5**. The p-values between EGPN and other eight GO prediction methods for $Fmax$ and $AUPR$ on 8 species.

**Table S6**. The p-values between EGN and other six GO prediction methods for $Fmax$ and

$AUPR$ on 98 non-coding genes.

**Table S7**. The details of training and test dataset for 7 species in CAFA3 dataset.

**Table S8**. The p-values between TNP and other five expression profile-based methods for $Fmax$ and $AUPR$ on 2433 proteins of 7 species from CAFA3 test dataset.

**Table S9**. The p-values between TNP and other five expression profile-based methods for $Fmax$ and $AUPR$ on CAFA3 test dataset for each of 7 species

**Table S10**. The p-values between TripletGO and other six GO prediction methods for $Fmax$ and $AUPR$ on 2433 proteins of 7 species from CAFA3 test dataset

**Table S11**. The numbers of genes with GO annotation of three aspects for 20 species.

**Table S12**. The details of 8 benchmark datasets constructed in our work.

**Table S13**. The values of $\alpha$, $h$, $margin$, and $c_f$ on the benchmark datasets for 8 species.

#### Supporting Figures

**Figure S1**: The performance of six expression profile-based methods on the test datasets for 8 species.

**Figure S2**: The precision-recall curves of six expression profile-based methods on the test datasets for 8 species.

**Figure S3**: Variation curves of $Fmax$ values of TNP and NON-PCA-TNP on the test dataset of human species versus the sampling ratios in expression data.

**Figure S4**: The scattering plots of mean expression level versus F1-scores for 1470 human test genes by TNP.

**Figure S5**: The $AVG\_WFS$ values of six measures for three GO aspects in 8 individual species.

**Figure S6**: The scattering plots of weights versus F1-scores of 100 templates for the gene MIRLET7C over TNP, MR, and PCC.

**Figure S7**: The $Fmax$ values of nine GO prediction methods on the test datasets for 8 species.

**Figure S8**: The $AUPR$ values of nine GO prediction methods on the test datasets for 8 species.

**Figure S9**: The precision-recall curves of five GO prediction methods on the test datasets for 8 species.

**Figure S10**: Comparison of mean and median AUROC values of three GO aspects by different methods on the common dataset.

**Figure S11**: Comparison of $Fmax$ and $AUPR$ values of three GO aspects by different methods on the common dataset.

**Figure S12**: The performance of six expression profile-based methods for 7 species on CAFA3 test dataset

**Figure S13**: The directed acyclic graph of predicted GO terms with corresponding confidence scores for gene GALNT4 by TripletGO

**Figure S14**: The distribution of sequence identities for 10000 gene-gene pairs and 10000 mapped protein-protein pairs.

### Supporting Texts

#### Text S1. The procedures of GBA strategy for expression profile-based GO prediction

In the guilty-by-association (GBA) strategy, we select the template genes which have the highest similarity with query gene in terms of expression profiles, and then use the GO terms of templates to annotate the query, as follows.

##### Training stage

In a training dataset, the expression profiles of all genes can be represented as a matrix $\boldsymbol{E} = (e_{ij})_{m \times l}$, where the $i$-th row of $\boldsymbol{E}$ is the expression profile for the $i$-th gene and denoted as $\boldsymbol{e}_i = (e_{i1}, e_{i2}, \dots, e_{il})^T$, $m$ is the total number of training genes, $l$ is the number of experimental samples in microarray technology [1], and $e_{ij}$ is the expression value of the $i$-th gene on the $j$-th sample. We orderly execute z-score normalization [2] and principal component analysis (PCA) [3] on expression profile matrix $\boldsymbol{E}$ to obtain a normalized matrix $\boldsymbol{E}^n = (e_{ij}^n)_{m \times h}$, where the $i$-th row of $\boldsymbol{E}^n$, denoted as $\boldsymbol{e}_i^n = (e_{i1}^n, e_{i2}^n, \dots, e_{ih}^n)^T$, is the normalized expression profile vector for the $i$-th training gene.

##### Prediction stage

For a query gene, its expression profile can be represented as a vector $\boldsymbol{e}^q = (e_1^q, e_2^q, \dots, e_l^q)^T$. First, the z-score normalization and PCA are orderly executed on the expression profile vector $\boldsymbol{e}^q$ to obtain a normalized vector $\boldsymbol{e}_q^n = \left(e_1^{nq}, e_2^{nq}, \dots, e_h^{nq}\right)^T$. Then, for each training gene $i$, we calculate its similarity score with query based on the normalized vector $\boldsymbol{e}_q^n$ and $\boldsymbol{e}_i^n$. Next, we rank $m$ training genes based on the similarity scores in descending order. Finally, we select the top $K$ training genes as templates to annotate the GO terms of query. Specifically, the confidence score that the query is associated with GO term $Q_j$ can be calculated as follows:

$$S(Q_j)_{GBA} = \frac{\sum_{k=1}^{K} w_k \cdot I_k(Q_j)}{\sum_{k=1}^{K} w_k} \quad \text{(S1)}$$

$$w_k = 1 - (r_k - 1)/K \quad \text{(S2)}$$

where $w_k$ is the weight for the $k$-th template, and $r_k$ is the rank of the $k$-th template; $I_k(Q_j) = 1$, if the $k$-th template is associated with $Q_j$ in the experimental annotation;

otherwise, $I_k(Q_j) = 0$.

In this work, the similarity score of expression profiles between two genes are measured by four unsupervised methods, including Pearson correlation coefficient (PCC) [4], Spearman rank correlation (SRC) [5], mutual rank (MR) [6], and Euclidean distance (ED) [7], and a recently proposed supervised method, i.e., metric learning for co-expression (MLC) [8].

The PCC between the $i$-th training gene and query gene is calculated as follows:

$$\mathrm{PCC}(\boldsymbol{e}_i^n, \boldsymbol{e}_q^n) = \frac{\sum_{j=1}^{h}(e_{ij}^n - \overline{\boldsymbol{e}_i^n})\cdot(e_j^{nq} - \overline{\boldsymbol{e}_q^n})}{\sqrt{\sum_{i=1}^{h}(e_{ij}^n - \overline{\boldsymbol{e}_i^n})^2}\cdot\sqrt{\sum_{i=1}^{h}(e_j^{nq} - \overline{\boldsymbol{e}_q^n})^2}} \tag{S3}$$

where $\overline{\boldsymbol{e}_i^n}$ and $\overline{\boldsymbol{e}_q^n}$ are mean values for $\boldsymbol{e}_i^n$ and $\boldsymbol{e}_q^n$, respectively.

The SRC between the $i$-th training gene and query gene is calculated as follows:

$$\mathrm{SRC}(\boldsymbol{e}_i^n, \boldsymbol{e}_q^n) = 1 - \frac{6\sum_{j=1}^{h}(r_{ij} - r_j)^2}{h(h^2-1)} \tag{S4}$$

where $r_{ij}$ is rank of $e_{ij}^n$ in the elements of $\boldsymbol{e}_i^n$ in ascending order, $r_j$ is the rank of $e_j^{nq}$ in the elements of $\boldsymbol{e}_q^n$ in ascending order.

Due to the long computation time of MR values, we directly download MR values of genes from COXPRESdb [9] and ATTED-II databases [6]. In a species with $M$ genes, the MR value between gene $i$ and gene $j$ is calculated as follows. First, we calculate the PCC values between gene $i$ and the remaining $M-1$ genes based on the corresponding expression profile vectors, and rank the $M-1$ genes based on the PCC values in descending order. Similarly, we calculate the PCC values between gene $j$ and the remaining $M-1$ genes, and rank the $M-1$ genes in descending order based on PCC values. Then, the MR value between genes $i$ and $j$ can be calculated:

$$MR(i,j) = \sqrt{rank(i)\cdot rank(j)} \tag{S5}$$

where $rank(i)$ is the rank of gene $i$ in $M-1$ genes for gene $j$, and $rank(j)$ is the rank of gene $j$ in $M-1$ genes for gene $i$.

The ED between the $i$-th training gene and query gene is calculated as follows:

$$\mathrm{ED}(\boldsymbol{e}_i^n, \boldsymbol{e}_q^n) = \sqrt{\sum_{j=1}^{h}\left(e_{ij}^n - e_j^{nq}\right)^2} \tag{S6}$$

In MLC, the similarity between the $i$-th training gene and query gene is measured by weight inner product (WIP) as follows:

$$\mathrm{WIP}(\boldsymbol{e}_i^n, \boldsymbol{e}_q^n) = (\boldsymbol{e}_i^n)^T\cdot W\cdot \boldsymbol{e}_q^n \tag{S7}$$

where $W = \mathrm{diag}(w)$ is a diagonal matrix and can be optimized by the Broyden-

Fletcher-Goldfarb-Shanno method [10].

The higher values of PCC, SRC, and WIP indicate the higher similarity, while the lower values of MR and ED mean the higher similarity.

**Text S2. The performances of six expression profile-based GO prediction methods for each individual species**

For each of 8 species, we will evaluate the performances of six expression profile-based GO prediction methods on the corresponding test dataset. For each method, we execute it 10 times and then use the average of all prediction results as the final result.

Figure S1 show the values of $Fmax$ and $AUPR$ for 8 species via six expression profile-based methods. Table S2 summarizes the p-values of $Fmax$ and $AUPR$ values between TNP and other five methods in student's t-test [11] for 8 species. In comparison between TNP and MLC, we use two samples t-test [12] to calculate p-value due to that the prediction results in 10 times are different for MLC/TNP. In comparison between TNP and PCC, MR, SRC, ED, we use single samples t-test [13] to calculate p-value, because the prediction results in 10 times are same for PCC/MR/SRC/ED. From Figure S1 and Table S2, we can observe that TNP achieves the highest values of $Fmax$ and $AUPR$ among six methods for each GO aspect in each species. For example, in human species, the improvements of $Fmax$ values between TNP and MR are 12.7%, 8.2%, 3.8%, respectively, with p-values of $1.29\times10^{-04}$, $7.83\times10^{-09}$, and $5.07\times10^{-07}$ for MF, BP, and CC aspects. As another example, the average improvement of $AUPR$ values of three GO aspects between TNP and the second best performer is 8.6% with p-values<0.05 for arabidopsis species.

Figure S2 plots the precision-recall (PR) curves of six expression profile-based methods for three GO aspects in 8 species. For each GO aspect in each species, we can find that TNP has the highest precision values among six expression profile-based methods at all different recall rates.

**Text S3. Exploring the influence of the characteristics of expression data on prediction performance for human species**

We used the human data to explore the impact of the characteristics of expression data on the prediction performance of TNP from the following two aspects.

First, to analyze the dependency of the prediction performance of TNP on the number of expression samples, we deigned the following test. In human benchmark dataset, each gene is associated with 27,655 expression samples, as shown in Table S11. For each human gene, we randomly extracted 10%, 30%, 50%, and 70% expression samples, respectively. This generates four sub-datasets from the original human benchmark set. Each sub-dataset has the same genes but different numbers (i.e., 2,765, 8,296, 13,827, and 19,358) of expression samples. In each sub-benchmark dataset, we re-trained the TNP model on the training dataset, and then evaluated the corresponding prediction performance on the test dataset. Moreover, to eliminate the redundant information of expression samples, TNP uses the principal component analysis (PCA) to reduce the dimension of expression profile vector. To further demonstrate the effectiveness of PCA, we added a comparison model, named NON-PCA-TNP, which removes the PCA from TNP.

Table S3 summarizes the $Fmax$ values of TNP and NON-PCA-TNP on the test dataset of human species for different sampling ratios in expression data. Figure S3 plots the variation curves of $Fmax$ values of two methods versus the sampling ratios. From Table S3 and Figure S3, the following two conclusions can be drawn. First, the performance of TNP has a low dependency on the number of expression samples. Specifically, with the increase of sampling ratio from 10% to 100%, the performance of TNP is slowly improved. Under the sampling ratio of 30%, TNP can achieve the acceptable $Fmax$ values, which are only decreased on average by 1.2% on three GO aspects in comparison with the model using all expression samples. Second, the PCA is helpful for relieving information redundancy among expression samples. Concretely, we found that the performance of TNP is consistently better than that of NON-PCA-TNP under all sampling ratios for each GO aspect.

In addition, to analyze the relation between prediction performance and expression level for individual gene, we further deigned the following test. For each test gene in human species, we firstly calculated the mean expression level of all expression samples and the F1-score between the predicted GO terms by TNP and actual GO terms in native annotation. Then, the Pearson correlation coefficient (PCC) between mean expression level and F1-score for all test genes was calculated.

Figure S4 plots the scattering plots of mean expression level versus F1-score for 1,470 human test genes by TNP, where the PCC value between mean expression level and F1-score is listed at the top. The PCC between mean expression level and F1-score ranges from -0.026 to 0.173. Therefore, we can conclude that there is no clear correlation between expression level and prediction performance of TNP for individual genes.

**Text S4. The functional similarity for genes**

The functional similarity of two genes is measured by the F1-score between their experimental GO terms. For a gene pair $(i, j)$, the F1-score between their GO terms is defined as:

$$\mathrm{F1-score} = 2(\mathrm{pre} \times \mathrm{rec})/(\mathrm{pre} + \mathrm{rec}),\ \mathrm{pre} = ns/n_1,\ \mathrm{rec} = ns/n_2 \quad \text{(S8)}$$

where $ns$ is the number of same GO terms between two genes, $n_1$ and $n_2$ are the numbers of GO terms for genes $i$ and $j$, respectively.

**Text S5. The performances of nine GO prediction methods for each individual species**

For each of 8 species, we will compare the performances of four individual methods (i.e., EPGP, GSAGP, PSAGP, and NGP) and five combination methods (i.e., GPN, EPN, EGN, EGP, and EGPN) on the corresponding test dataset. For each combination method, we execute it 10 times and then use the average of all prediction results as the final result.

Figures S7-S8 illustrate the values of $Fmax$ and $AUPR$ for nine GO prediction methods in 8 species. Table S5 show the p-values of $Fmax$ and $AUPR$ values between EGPN and other eight methods in student's t-test [11] for 8 species. In comparison between EGPN and four combination methods (i.e., GPN, EPN, EGN, and EGP), we use two samples t-test [12] to calculate p-value due to that the prediction results in 10 times are different for them. In comparison between EGPN and four individual methods (i.e., EPGP, GSAGP, PSAGP, and NGP), we use single samples t-test [13] to calculate p-value. From Figures S7-S8 and Table S5, we can find the $Fmax$ and $AUPR$ values of EGPN are much higher than that of four individual methods for each species. Moreover, from the view of $Fmax$, in species of arabidopsis and fly, EGPN achieves the better performance than other four combination methods for each GO aspect; in species of human, mouse, rat and nematoda, EGPN occupies one of the top two positions among five combination methods for each GO aspect; as for the remaining two species, EGPN shows the best performance in MF/BP for budding yeast and BP/CC for fission yeast. These observations further demonstrate that each individual method contributes to improving prediction performance.

Figure S9 plots the precision-recall (PR) curves of four individual methods and EPGN for three GO aspects in 8 species. For each GO aspect in each species, we can observe that the PR curve of EGPN is continuously higher than that of four individual methods.

**Text S6. Finding common genes and GO terms between our datasets and GENETICA's datasets**

In the web page (http://genetica-network.com), GENETICA provides the prediction scores and real labels of three GO aspects for 19,635 genes in human species and 18425 genes in mouse species. Specifically, in human species, each gene is associated with the prediction scores and real labels of 843 MF, 4203 BP, and 528 CC GO terms. As for mouse species, each gene is associated with the prediction scores and real labels of 833 MF, 4188 BP, and 525 CC terms.

In GENETICA's datasets, 879 genes, 1176 genes, and 1241 genes for MF, BP, and CC aspects, respectively, can be found in our test dataset for human species; 572 genes, 880 genes, and 737 genes for three GO aspects can be separately found in our test dataset for mouse species. The GO terms in GENETICA and our work are represented as GO names and GO IDs, respectively. Due to the different versions of gene ontology databases, only 738 MF, 3980 BP and 476 CC GO names in GENETICA's can be correctly mapped as the corresponding GO IDs in our dataset for human species; As for mouse species, there are 727 MF, 3965 BP, and 474 CC terms in common between our work and GENETICA. Moreover, we only consider the GO terms whose frequencies are more than 20 both in our training datasets and GENETICA's datasets. After this, there are 879 genes annotated with 287 MF terms, 1176 genes with 1340 BP terms, and 1241 genes with 186 CC terms for human species in common between our test dataset and GENETICA's dataset; As for mouse species, there are 572 genes with 149 MF terms, 880 genes with 1230 BP terms, and 737 genes with 128 CC terms in common. For each GO term $Q_i$, all genes are assigned with the prediction scores and real labels. Specifically, if a gene is associated with $Q_i$ both in our test dataset and GENETICA's dataset, we label it as "1"; otherwise, it is labeled as "0".

**Text S7. Finding common genes and GO terms between our datasets and GeneNetwork's datasets**

In the web page (https://www.genenetwork.nl/), we can use command "GET https://www.genenetwork.nl/api/v1/gene/geneName?db=database” to download the GO information file generated by GeneNetwork for each query gene in each GO aspect. The information file contains all GO terms in the experimental function annotation and 100 predicted GO terms with scores for a query. GeneNetwork provides all of the information files in three GO aspects for 56435 genes.

In GeneNetwork's dataset, 918 genes, 1230 genes, and 1328 genes for MF, BP and CC aspects, respectively, can be found in our test dataset for human species. Moreover, there are 655 MF, 2776 BP, and 536 CC terms in common between our work and GeneNetwork. In this work, we only consider the GO terms whose frequencies are more than 20 both in our training dataset and GeneNetwork's dataset. After this, there are 918 genes associated with 165 MF terms, 1230 genes with 522 BP terms, and 1328 genes with 182 CC terms, in common, for human species between our test dataset and GeneNetwork's dataset. For each GO term $Q_i$, all genes are assigned with the prediction scores and real labels. Specifically, if a gene is associated with $Q_i$ both in our test dataset and GeneNetwork's dataset, we label it as “1”; otherwise, it is labeled as “0”.

**Text S8. Comparison with the existing gene function prediction models in gene-center level.**

We further compared our methods (TNP and TripletGO) with the existing gene function predictors (GENETICA and GeneNetwork) in gene-center level. In GENETICA and GeneNetwork, the numerical distributions of prediction scores of genes are different for each GO aspect. For example, in MF aspect, the prediction scores of GO terms for all human genes by GENETICA are range from 0.618167 to 15.79360; in BP aspect, the prediction scores by GENETICA are range from 0.61976 to 19.75050. Therefore, we firstly normalize the prediction scores in the range of 0 to 1 for each GO aspect using Min-Max Normalization. Specifically, for each GO aspect, the prediction score $S_i$ is normalized as:

$$S_i^n = \frac{S_i - S_{min}}{S_{max} - S_{min}} \quad \text{(S9)}$$

where $S_{max}$ and $S_{min}$ are the max and min values, respectively, in all prediction scores.

Based on the normalized prediction scores, the prediction performances for GENETICA and GeneNetwork can be transformed from term-center metric (AUROC) to gene-center metric ($Fmax$ and $AUPR$). Figure S11 (A-B) shows the $Fmax$ and $AUPR$ values of TNP, TripletGO and GENETICA in human species (879 genes with 287 MF terms, 1176 genes with 1340 BP terms, and 1241 genes with 186 CC terms) and mouse species (572 genes with 149 MF terms, 880 genes with 1230 BP terms, and 737 genes with 128 CC terms). Moreover, we further compared our methods with GeneNetwork in human species (918 genes with 165 MF terms, 1230 genes with 522 BP terms, and 1328 genes with 182 CC terms), as shown in Figure S11 (C). From Figure S11 (A-C), we find that the proposed TNP and TripletGO show the significantly better performance than GENETICA and GeneNetwork for each GO aspect. For example, from the view of $Fmax$, TNP achieves the improvements of 112.8%, 60.7% and 59.1%, respectively, for MF, BP, and CC aspects of human species in comparison with GENETICA. It cannot escape our notice that the $Fmax$ and $AUPR$ values of TNP and TripletGO in Figure S11 are significantly lower than the corresponding values in the previous figures (Figures S1, S7 and S8), especially for CC aspect. The reason can be explained as follows. Frist, we remove a part of genes and terms, which are not included in the GENETICA's dataset, from our test dataset. Second, the GO annotations of genes for our work and GENETICA are originated from different databases. Specifically, the GO annotation of genes in our work are downloaded from NCBI with the version of "2021-02-23"; in GENETICA, the GO annotations are extracted from Broad Institute Molecular Signatures Database v6.2. Therefore, the GO annotations for a part of test genes are different between our work and GENETICA. For example, for gene ARHGAP1 (Entrez ID: 392), we listed the corresponding GO annotations for CC aspect in different works as follows.

(1) Our work: GO:0005768, GO:0110165, GO:0016020, GO:0043227, GO:0043226, GO:0097708, GO:0005737, GO:0031982, GO:0097443, GO:0031410, GO:0010008, GO:0098588, GO:0031090, GO:0048471, GO:0005829 (15 terms).

(2) GENETICA: GO:0048471, GO:0110165 (2 terms)

(3) GeneNetwork: GO:0005737, GO:0016020, GO:0043230, GO:0070062, GO:0005829, GO:0098588, GO:0010008, GO:0031090, GO:0048471, GO:1903561, GO:0043227, GO:0043226, GO:0110165, GO:0031982 (14 terms)

(4) Common terms between our work and GENETICA: GO:0048471, GO:0110165 (2 terms)

(5) Common terms between our work and GeneNetwork: GO:0005737, GO:0031090, GO:0005829, GO:0016020, GO:0031982, GO:0043226, GO:0048471, GO:0043227, GO:0098588, GO:0110165, GO:0010008 (11 terms).

We can notice that there are only 2 common GO terms (GO:0048471, GO:0110165) for the annotation of gene ARHGAP1 between our work and GENETICA. To compare our method and GENETICA in fairness, these 2 common GO terms are used as "gold standard" for GO annotation of gene ARHGAP1 and the remaining terms are ignored. Therefore, some predicted terms, which are considered as true positives in previous Figures (Figures S1, S7 and S8), are viewed as false positives in the comparison with GENETICA, which further leads the significant performance degradation of TNP and TripletGO in this section.

**Text S9. The performances of six expression profile-based GO prediction methods for each individual species on CAFA3 test dataset**

We further benchmarked the performances of six expression profile-based GO prediction methods for each of 7 species on CAFA3 test dataset. Figure S12 shows the values of $Fmax$ and $AUPR$ for 7 species via six expression profile-based methods. Table S9 summarizes the p-values of $Fmax$ and $AUPR$ values between TNP and other five methods in student's t-test [11] for 7 species. We can observe that TNP achieves the highest values of $Fmax$ and $AUPR$ among six methods for each GO aspect in most of 7 species. Taking human species as an example, TNP achieves 11.9% and 7.8% average increases on three GO aspects for $Fmax$ and $AUPR$, respectively, in comparison with the second-best performer, i.e., MR.

**Text S10. The confidence scores of the candidate GO terms for gene GALNT4 by TripletGO**

Cut-off value: 0.350

Confidence scores of the candidate GO terms:

**GO:0110165 C 0.968 GO:0016020 C 0.771 GO:0031090 C 0.700 GO:0043226 C 0.626 GO:0043227 C 0.602 GO:0098588 C 0.534 GO:0005622 C 0.417 GO:0043229 C 0.415 GO:0000139 C 0.405 GO:0043231 C 0.369**
GO:0031982 C 0.325 GO:0097708 C 0.284 GO:0031410 C 0.284 GO:0005886 C 0.265 GO:0005829 C 0.184 GO:0005789 C 0.177 GO:0030133 C 0.166 GO:0005654 C 0.141 GO:0005737 C 0.136 GO:0005783 C 0.135 GO:0031224 C 0.134 GO:0016021 C 0.134 GO:0005794 C 0.112 GO:0032991 C 0.087 GO:0031226 C 0.084 GO:0005887 C 0.084 GO:0005634 C 0.070 GO:0043228 C 0.049 GO:0043232 C 0.047 GO:0098590 C 0.045 GO:0098796 C 0.043 GO:0031974 C 0.037 GO:0012506 C 0.037 GO:0042995 C 0.036 GO:0120025 C 0.036 GO:0043233 C 0.035 GO:0030659 C 0.034 GO:0005768 C 0.031 GO:0070013 C 0.027 GO:0030054 C 0.024 GO:0005739 C 0.022 GO:0031301 C 0.022 GO:0031300 C 0.022 GO:0010008 C 0.021 GO:1902494 C 0.019 GO:0048471 C 0.019 GO:0030667 C 0.018 GO:0016604 C 0.017 GO:0005815 C 0.015 GO:0005576 C 0.014 GO:0005774 C 0.012 GO:0098852 C 0.012 GO:0005765 C 0.012 GO:1990234 C 0.011 GO:0070161 C 0.010 GO:0005929 C 0.010

**Text S11. The construction procedures of Gene-GOA**

First, we download all genes with GO annotation from National Center for Biotechnology Information [14] (NCBI). Following the CAFA experiments [15, 16], we only select the genes annotated by at least one of the eight experimental evidence codes, including EXP, IDA, IPI, IMP, IGI, IEP, TAS, and IC. Moreover, to explicitly consider the hierarchical structure of GO terms, if a child term is annotated to a gene, all its direct and indirect parents, as defined by the "is_a" relation in gene ontology database [17] (http://geneontology.org/), are also annotated. The numbers of genes annotated with GO terms for MF, BP, and CC are 40160, 63543, and 55448, respectively, in Gene-GOA.

**Text S12. Distance rank-based strategy**

The distance rank-based strategy (DRBS) is executed on normalized embedding matrix of training genes ($\boldsymbol{U}^n$) and normalized embedding vector of query gene ($\boldsymbol{u}^q$) to obtain a confidence score vector, denoted as $\boldsymbol{s}^t = (s_1^t, s_2^t, \dots, s_r^t)^T$, where $s_i^t$ is the confidence score that query is associated with the $i$-th GO term from the view of distance rank in the embedding space.

The details of DRBS are described as follows. First, we rank $m$ training genes based the distances between the training and query genes in embedding space in ascending order. The distance between the $i$-th training gene and query gene is calculated as follows:

$$d(i, query) = \sum_{k=1}^{d_N} \left(u_{ik}^n - u_k^q\right)^2 / 4 \tag{S10}$$

Then, we select top $K$ training genes which have the shortest distance with query in embedding space as templates to calculate the confidence scores of GO terms for query as follows:

$$s_i^t = \frac{\sum_{k=1}^{K} w_k \cdot I_k(i)}{\sum_{k=1}^{K} w_k} \tag{S11}$$

$$w_k = 1 - (r_k - 1)/K \tag{S12}$$

where $w_k$ is the weight for the $k$-th template, and $r_k$ is the rank of the $k$-th template; $I_k(i) = 1$, if the $k$-th template is associated with the $j$-th GO term in the experimental function annotation; otherwise, $I_k(i) = 0$. In this work, the value of $K$ is set to be 100.

**Text S13. The construction procedures of genetic sequence database with GO annotation**

To construct a genetic sequence database with GO annotation (GSD-GOA), the RNA sequences of all genes in Gene-GOA are extracted from NCBI [14]. If there is no available RNA sequence for a gene, its genomic DNA sequence is selected. In addition, we discard a few genes which have no available RNA or genomic DNA sequences in NCBI. After this, GSD-GOA includes 39179 sequences with MF terms, 61699 sequences with BP terms, and 54117 sequences with CC terms.

**Text S14. The construction procedures of protein sequence database**

The protein sequence database (PSD) is constructed as follows. For each gene in Gene-GOA, we map it as the corresponding coding protein sequences in UniProt database [18] using a gene-protein mapping table. After this, 78170 genes can be mapped as 119876 protein sequences.

**Text S15. The relationship between protein sequence identity and genetic sequence identity**

In this work, we use three machine learning models, including liner regression (LR) [19], support vector regression (SVR) [20], and neural network with one hidden layer (NN), to fit the relationship between protein sequence identity and genetic sequence identity, as follows:

$$y = f(x)_{\mathrm{Model}} \quad \text{(S13)}$$

where $x$ is the protein sequence identity, $y$ is the genetic sequence identity, $\mathrm{Model} \in \{\mathrm{LR}, \mathrm{SVR}, \mathrm{NN}\}$.

First, we randomly select 100,000,000 gene-gene sequence pairs from NCBI, and map each gene-gene sequence pair as a protein-protein sequence pair in UniProt database by gene-protein mapping table. Then, to reduce computation time, we remove the protein sequences and gene sequences whose lengths are more than 10000. After this, the number of remaining gene-gene pairs (or the corresponding protein-protein pairs) is 974447. Next, we use standard Needleman-Wunsch algorithm [21] to calculate the sequence identity for each gene-gene pair and the corresponding protein-protein pair. Each gene-gene pair and the corresponding protein-protein pair are combined as a machine learning sample. More specifically, the sequence identity of protein-protein pair is used as the feature of sample, i.e., the input of machine learning model, and the sequence identity of gene-gene pair is used as the regression value of sample, i.e., the output of machine learning model. Finally, we separately train LR, SVR and NN on the 974447 samples, and found that $f(30\%)_{LR} = 58.6\%$, $f(30\%)_{SVM} = 59.3\%$, $f(30\%)_{NN} = 60.1\%$. In addition, we randomly select 10,000 gene-gene pairs and the mapped protein-protein pairs, and then plot the corresponding sequence identities, as shown in Figure S14. For the protein-protein pairs whose sequence identities are equal to 30%, the sequence identities for the corresponding gene-gene pairs are near to 60%. In light of the above, we use 60% sequence identity as the cut-off to remove the homologous gene templates.

### Supporting Tables

**Table S1 The p-values between TNP and other five expression profile-based methods for $WAFmax$ and $WAAUPR$**

| Measure | GO aspect | (TNP, MR) | (TNP, PCC) | (TNP, MLC) | (TNP, SRC) | (TNP, ED) |
|---|---|---|---|---|---|---|
| $WAFmax$ | MF | $8.60\times10^{-08}$ | $1.70\times10^{-10}$ | $6.56\times10^{-09}$ | $3.19\times10^{-09}$ | $8.25\times10^{-11}$ |
| | BP | $6.11\times10^{-11}$ | $1.33\times10^{-13}$ | $1.71\times10^{-12}$ | $5.65\times10^{-13}$ | $8.64\times10^{-14}$ |
| | CC | $9.24\times10^{-09}$ | $3.67\times10^{-11}$ | $1.21\times10^{-09}$ | $3.85\times10^{-10}$ | $5.60\times10^{-11}$ |
| $WAAUPR$ | MF | $5.13\times10^{-10}$ | $1.39\times10^{-14}$ | $1.14\times10^{-11}$ | $3.25\times10^{-13}$ | $1.04\times10^{-14}$ |
| | BP | $3.04\times10^{-09}$ | $4.43\times10^{-13}$ | $7.37\times10^{-12}$ | $2.61\times10^{-12}$ | $6.69\times10^{-13}$ |
| | CC | $1.82\times10^{-13}$ | $3.54\times10^{-15}$ | $1.00\times10^{-13}$ | $1.00\times10^{-13}$ | $6.58\times10^{-15}$ |

**Table S2 The p-values between TNP and other five expression profile-based methods for $Fmax$ and $AUPR$ on 8 species**

| Species | Measure | GO aspect | (TNP, MR) | (TNP, PCC) | (TNP, MLC) | (TNP, SRC) | (TNP, ED) |
|---|---|---|---|---|---|---|---|
| Human | $Fmax$ | MF | $1.29\times10^{-04}$ | $3.08\times10^{-08}$ | $2.09\times10^{-12}$ | $1.80\times10^{-05}$ | $7.39\times10^{-08}$ |
| | | BP | $7.83\times10^{-09}$ | $2.32\times10^{-11}$ | $7.00\times10^{-18}$ | $2.00\times10^{-10}$ | $3.23\times10^{-11}$ |
| | | CC | $5.07\times10^{-07}$ | $1.95\times10^{-10}$ | $8.01\times10^{-09}$ | $2.53\times10^{-09}$ | $4.75\times10^{-10}$ |
| | $AUPR$ | MF | $2.42\times10^{-07}$ | $3.72\times10^{-11}$ | $1.87\times10^{-16}$ | $7.15\times10^{-09}$ | $2.95\times10^{-11}$ |
| | | BP | $1.42\times10^{-07}$ | $1.51\times10^{-11}$ | $1.11\times10^{-16}$ | $6.09\times10^{-10}$ | $3.94\times10^{-11}$ |
| | | CC | $4.64\times10^{-12}$ | $4.30\times10^{-13}$ | $5.30\times10^{-17}$ | $4.64\times10^{-12}$ | $1.52\times10^{-12}$ |
| Mouse | $Fmax$ | MF | $2.58\times10^{-04}$ | $4.38\times10^{-07}$ | $1.89\times10^{-10}$ | $4.09\times10^{-03}$ | $4.00\times10^{-09}$ |
| | | BP | $1.24\times10^{-03}$ | $3.46\times10^{-09}$ | $1.16\times10^{-13}$ | $1.48\times10^{-04}$ | $4.55\times10^{-09}$ |
| | | CC | $2.96\times10^{-05}$ | $5.33\times10^{-07}$ | $1.22\times10^{-12}$ | $4.56\times10^{-06}$ | $3.73\times10^{-07}$ |
| | $AUPR$ | MF | $2.01\times10^{-06}$ | $1.51\times10^{-10}$ | $2.07\times10^{-14}$ | $4.76\times10^{-06}$ | $4.98\times10^{-11}$ |
| | | BP | $7.61\times10^{-04}$ | $5.81\times10^{-09}$ | $1.48\times10^{-14}$ | $4.25\times10^{-06}$ | $5.81\times10^{-09}$ |
| | | CC | $1.65\times10^{-04}$ | $2.14\times10^{-08}$ | $5.98\times10^{-15}$ | $1.65\times10^{-04}$ | $2.91\times10^{-09}$ |
| Arabidopsis | $Fmax$ | MF | $1.74\times10^{-08}$ | $5.57\times10^{-10}$ | $3.27\times10^{-16}$ | $1.76\times10^{-09}$ | $1.17\times10^{-09}$ |
| | | BP | $2.91\times10^{-07}$ | $2.17\times10^{-09}$ | $7.00\times10^{-12}$ | $1.27\times10^{-07}$ | $6.07\times10^{-10}$ |
| | | CC | $7.50\times10^{-09}$ | $2.30\times10^{-10}$ | $1.07\times10^{-09}$ | $2.27\times10^{-06}$ | $5.81\times10^{-10}$ |
| | $AUPR$ | MF | $2.89\times10^{-07}$ | $3.47\times10^{-11}$ | $2.84\times10^{-14}$ | $1.73\times10^{-07}$ | $7.94\times10^{-11}$ |
| | | BP | $1.76\times10^{-06}$ | $6.41\times10^{-10}$ | $4.53\times10^{-13}$ | $1.37\times10^{-07}$ | $1.22\times10^{-09}$ |
| | | CC | $1.82\times10^{-11}$ | $1.39\times10^{-12}$ | $3.84\times10^{-15}$ | $6.67\times10^{-10}$ | $5.29\times10^{-12}$ |
| Rat | $Fmax$ | MF | $3.04\times10^{-01}$ | $6.57\times10^{-04}$ | $9.63\times10^{-07}$ | $1.87\times10^{-01}$ | $1.43\times10^{-04}$ |
| | | BP | $1.49\times10^{-03}$ | $4.76\times10^{-05}$ | $1.31\times10^{-08}$ | $1.02\times10^{-02}$ | $1.09\times10^{-05}$ |
| | | CC | $2.04\times10^{-01}$ | $2.30\times10^{-04}$ | $9.75\times10^{-07}$ | $9.68\times10^{-03}$ | $2.01\times10^{-04}$ |
| | $AUPR$ | MF | $9.02\times10^{-04}$ | $3.74\times10^{-05}$ | $1.43\times10^{-05}$ | $8.78\times10^{-06}$ | $1.63\times10^{-05}$ |
| | | BP | $1.40\times10^{-02}$ | $3.60\times10^{-06}$ | $4.04\times10^{-10}$ | $1.46\times10^{-01}$ | $1.30\times10^{-05}$ |
| | | CC | $9.00\times10^{-07}$ | $1.44\times10^{-10}$ | $1.17\times10^{-08}$ | $1.38\times10^{-07}$ | $6.65\times10^{-11}$ |
| Fly | $Fmax$ | MF | $6.69\times10^{-03}$ | $5.35\times10^{-06}$ | $9.31\times10^{-11}$ | $1.02\times10^{-07}$ | $3.97\times10^{-07}$ |
| | | BP | $7.87\times10^{-06}$ | $6.97\times10^{-09}$ | $5.39\times10^{-12}$ | $4.91\times10^{-10}$ | $1.97\times10^{-09}$ |
| | | CC | $3.01\times10^{-06}$ | $8.38\times10^{-07}$ | $2.77\times10^{-14}$ | $1.35\times10^{-10}$ | $5.11\times10^{-07}$ |
| | $AUPR$ | MF | $3.31\times10^{-01}$ | $1.55\times10^{-05}$ | $9.90\times10^{-14}$ | $6.06\times10^{-10}$ | $6.78\times10^{-06}$ |
| | | BP | $4.03\times10^{-06}$ | $6.66\times10^{-10}$ | $4.77\times10^{-13}$ | $1.34\times10^{-11}$ | $3.24\times10^{-10}$ |
| | | CC | $2.04\times10^{-08}$ | $2.56\times10^{-11}$ | $7.29\times10^{-15}$ | $3.58\times10^{-13}$ | $3.63\times10^{-12}$ |
| Budding Yeast | $Fmax$ | MF | $4.17\times10^{-06}$ | $1.05\times10^{-08}$ | $5.22\times10^{-13}$ | $1.05\times10^{-08}$ | $1.22\times10^{-08}$ |
| | | BP | $4.32\times10^{-06}$ | $7.34\times10^{-08}$ | $1.20\times10^{-13}$ | $2.07\times10^{-08}$ | $1.41\times10^{-08}$ |
| | | CC | $3.80\times10^{-05}$ | $4.07\times10^{-08}$ | $1.41\times10^{-11}$ | $1.65\times10^{-06}$ | $2.72\times10^{-08}$ |
| | $AUPR$ | MF | $9.24\times10^{-06}$ | $2.72\times10^{-09}$ | $2.75\times10^{-15}$ | $1.03\times10^{-10}$ | $2.13\times10^{-09}$ |
| | | BP | $2.98\times10^{-08}$ | $6.08\times10^{-11}$ | $2.64\times10^{-15}$ | $4.84\times10^{-12}$ | $7.64\times10^{-11}$ |
| | | CC | $8.16\times10^{-05}$ | $4.98\times10^{-08}$ | $8.94\times10^{-12}$ | $1.13\times10^{-08}$ | $2.91\times10^{-07}$ |
| Fission Yeast | $Fmax$ | MF | $1.47\times10^{-05}$ | $2.70\times10^{-02}$ | $4.26\times10^{-07}$ | $8.92\times10^{-04}$ | $2.70\times10^{-02}$ |
| | | BP | $1.05\times10^{-06}$ | $4.87\times10^{-06}$ | $6.31\times10^{-13}$ | $4.48\times10^{-08}$ | $4.87\times10^{-06}$ |
| | | CC | $9.97\times10^{-01}$ | $3.26\times10^{-01}$ | $8.52\times10^{-04}$ | $3.50\times10^{-04}$ | $3.26\times10^{-01}$ |
| | $AUPR$ | MF | $6.16\times10^{-03}$ | $8.69\times10^{-01}$ | $3.91\times10^{-08}$ | $8.36\times10^{-04}$ | $8.69\times10^{-01}$ |
| | | BP | $1.99\times10^{-06}$ | $6.24\times10^{-06}$ | $4.60\times10^{-12}$ | $6.24\times10^{-06}$ | $6.24\times10^{-06}$ |
| | | CC | $2.41\times10^{-03}$ | $2.63\times10^{-05}$ | $4.24\times10^{-07}$ | $3.51\times10^{-06}$ | $2.63\times10^{-05}$ |
| Nematoda | $Fmax$ | MF | $5.71\times10^{-04}$ | $5.20\times10^{-03}$ | $3.19\times10^{-08}$ | $2.68\times10^{-06}$ | $9.79\times10^{-04}$ |
| | | BP | $2.12\times10^{-06}$ | $9.91\times10^{-07}$ | $1.09\times10^{-10}$ | $4.80\times10^{-09}$ | $1.58\times10^{-07}$ |
| | | CC | $1.45\times10^{-02}$ | $1.22\times10^{-02}$ | $3.95\times10^{-10}$ | $6.02\times10^{-07}$ | $2.68\times10^{-03}$ |
| | $AUPR$ | MF | $5.09\times10^{-04}$ | $8.31\times10^{-05}$ | $1.83\times10^{-11}$ | $2.19\times10^{-08}$ | $1.10\times10^{-05}$ |
| | | BP | $2.09\times10^{-06}$ | $6.80\times10^{-09}$ | $2.96\times10^{-13}$ | $5.79\times10^{-12}$ | $3.42\times10^{-09}$ |
| | | CC | $3.90\times10^{-05}$ | $2.28\times10^{-07}$ | $6.39\times10^{-12}$ | $3.74\times10^{-08}$ | $2.28\times10^{-07}$ |

**Table S3 The $Fmax$ values of TNP and NON-PCA-TNP on the test dataset of human species for different sampling ratios in expression data**

| Method | GO aspect | 10% | 30% | 50% | 70% | 100% |
|---|---|---|---|---|---|---|
| TNP | MF | 0.293 | 0.307 | 0.311 | 0.307 | 0.311 |
| | BP | 0.388 | 0.391 | 0.393 | 0.392 | 0.397 |
| | CC | 0.565 | 0.573 | 0.573 | 0.574 | 0.577 |
| NON-PCA-TNP | MF | 0.287 | 0.291 | 0.292 | 0.294 | 0.290 |
| | BP | 0.382 | 0.384 | 0.385 | 0.386 | 0.385 |
| | CC | 0.564 | 0.571 | 0.572 | 0.572 | 0.573 |

**Table S4 The p-values between EGPN and other eight GO prediction methods for $WAFmax$ and $WAAUPR$ on the test datasets of 8 species**

| Measure | GO aspect | (EGPN, EPGP) | (EGPN, GSAGP) | (EGPN, PSAGP) | (EGPN, NGP) | (EGPN, GPN) | (EGPN, EPN) | (EGPN, EGN) | (EGPN, EGP) |
|---|---|---|---|---|---|---|---|---|---|
| $WAFmax$ | MF | $2.71\times10^{-21}$ | $3.85\times10^{-18}$ | $2.09\times10^{-14}$ | $9.57\times10^{-23}$ | $2.53\times10^{-10}$ | $7.12\times10^{-13}$ | $4.59\times10^{-17}$ | $1.70\times10^{-04}$ |
| | BP | $3.39\times10^{-15}$ | $4.82\times10^{-17}$ | $2.65\times10^{-15}$ | $1.42\times10^{-18}$ | $9.68\times10^{-13}$ | $6.33\times10^{-10}$ | $4.37\times10^{-12}$ | $4.34\times10^{-06}$ |
| | CC | $1.71\times10^{-13}$ | $7.18\times10^{-19}$ | $7.49\times10^{-18}$ | $2.07\times10^{-18}$ | $2.09\times10^{-14}$ | $8.98\times10^{-10}$ | $5.02\times10^{-11}$ | $1.96\times10^{-07}$ |
| $WAAUPR$ | MF | $3.31\times10^{-24}$ | $3.51\times10^{-24}$ | $1.17\times10^{-22}$ | $1.16\times10^{-25}$ | $3.09\times10^{-15}$ | $3.79\times10^{-17}$ | $2.31\times10^{-20}$ | $3.07\times10^{-04}$ |
| | BP | $6.93\times10^{-20}$ | $1.24\times10^{-23}$ | $2.99\times10^{-22}$ | $2.12\times10^{-23}$ | $3.66\times10^{-18}$ | $1.52\times10^{-14}$ | $4.97\times10^{-17}$ | $4.87\times10^{-11}$ |
| | CC | $9.69\times10^{-08}$ | $3.81\times10^{-16}$ | $3.32\times10^{-15}$ | $1.10\times10^{-14}$ | $2.70\times10^{-10}$ | $1.79\times10^{-05}$ | $2.73\times10^{-06}$ | $2.77\times10^{-04}$ |

**Table S5 The p-values between EGPN and other eight GO prediction methods for $Fmax$ and $AUPR$ on 8 species**

| Species | Measure | GO aspect | (EGPN, EPGP) | (EGPN, GSAGP) | (EGPN, PSAGP) | (EGPN, NGP) | (EGPN, GPN) | (EGPN, EPN) | (EGPN, EGN) | (EGPN, EGP) |
|---|---|---|---|---|---|---|---|---|---|---|
| Human | $Fmax$ | MF | $4.29\times10^{-16}$ | $4.45\times10^{-14}$ | $4.23\times10^{-08}$ | $1.37\times10^{-17}$ | $1.58\times10^{-03}$ | $1.26\times10^{-08}$ | $1.18\times10^{-21}$ | $2.65\times10^{-03}$ |
| | | BP | $2.04\times10^{-15}$ | $1.31\times10^{-17}$ | $5.53\times10^{-15}$ | $1.75\times10^{-18}$ | $7.79\times10^{-20}$ | $2.20\times10^{-11}$ | $2.76\times10^{-19}$ | $5.95\times10^{-11}$ |
| | | CC | $9.71\times10^{-12}$ | $1.57\times10^{-16}$ | $1.58\times10^{-15}$ | $8.73\times10^{-17}$ | $3.91\times10^{-12}$ | $7.89\times10^{-09}$ | $9.87\times10^{-14}$ | $6.36\times10^{-11}$ |
| | $AUPR$ | MF | $1.18\times10^{-21}$ | $2.03\times10^{-21}$ | $1.84\times10^{-19}$ | $4.08\times10^{-23}$ | $9.18\times10^{-18}$ | $7.82\times10^{-23}$ | $1.05\times10^{-31}$ | $3.66\times10^{-07}$ |
| | | BP | $2.48\times10^{-18}$ | $8.91\times10^{-22}$ | $2.70\times10^{-20}$ | $1.52\times10^{-21}$ | $1.18\times10^{-26}$ | $1.42\times10^{-17}$ | $8.86\times10^{-25}$ | $1.68\times10^{-15}$ |
| | | CC | $5.75\times10^{-04}$ | $3.72\times10^{-12}$ | $1.71\times10^{-11}$ | $3.28\times10^{-11}$ | $5.69\times10^{-08}$ | $4.07\times10^{-03}$ | $6.30\times10^{-03}$ | $5.40\times10^{-01}$ |
| Mouse | $Fmax$ | MF | $5.63\times10^{-21}$ | $3.45\times10^{-15}$ | $8.18\times10^{-12}$ | $3.72\times10^{-22}$ | $1.36\times10^{-04}$ | $2.08\times10^{-17}$ | $3.99\times10^{-23}$ | $1.42\times10^{-02}$ |
| | | BP | $3.56\times10^{-16}$ | $2.99\times10^{-16}$ | $1.96\times10^{-16}$ | $6.38\times10^{-19}$ | $1.74\times10^{-12}$ | $1.26\times10^{-12}$ | $3.48\times10^{-16}$ | $6.33\times10^{-06}$ |
| | | CC | $1.06\times10^{-12}$ | $1.69\times10^{-16}$ | $8.19\times10^{-17}$ | $1.79\times10^{-17}$ | $2.70\times10^{-18}$ | $1.56\times10^{-12}$ | $1.98\times10^{-10}$ | $6.37\times10^{-01}$ |
| | $AUPR$ | MF | $3.98\times10^{-22}$ | $2.04\times10^{-20}$ | $1.73\times10^{-19}$ | $4.01\times10^{-23}$ | $2.16\times10^{-09}$ | $1.19\times10^{-25}$ | $5.99\times10^{-28}$ | $3.62\times10^{-01}$ |
| | | BP | $2.60\times10^{-19}$ | $9.96\times10^{-22}$ | $1.70\times10^{-21}$ | $2.96\times10^{-22}$ | $3.74\times10^{-22}$ | $1.12\times10^{-22}$ | $2.83\times10^{-23}$ | $3.21\times10^{-11}$ |
| | | CC | $7.71\times10^{-06}$ | $9.57\times10^{-14}$ | $2.25\times10^{-13}$ | $3.96\times10^{-13}$ | $1.28\times10^{-11}$ | $1.43\times10^{-03}$ | $2.78\times10^{-03}$ | $2.79\times10^{-01}$ |
| Arabidopsis | $Fmax$ | MF | $1.52\times10^{-17}$ | $5.58\times10^{-14}$ | $1.06\times10^{-12}$ | $4.93\times10^{-19}$ | $1.39\times10^{-11}$ | $7.77\times10^{-15}$ | $7.09\times10^{-17}$ | $2.28\times10^{-03}$ |
| | | BP | $3.80\times10^{-11}$ | $1.67\times10^{-10}$ | $5.28\times10^{-11}$ | $1.01\times10^{-14}$ | $7.19\times10^{-09}$ | $1.04\times10^{-07}$ | $1.39\times10^{-07}$ | $8.80\times10^{-02}$ |
| | | CC | $1.58\times10^{-07}$ | $5.08\times10^{-14}$ | $1.80\times10^{-14}$ | $1.02\times10^{-12}$ | $3.19\times10^{-13}$ | $5.82\times10^{-08}$ | $2.30\times10^{-03}$ | $4.44\times10^{-02}$ |
| | $AUPR$ | MF | $1.07\times10^{-20}$ | $1.21\times10^{-21}$ | $2.38\times10^{-20}$ | $1.78\times10^{-22}$ | $1.15\times10^{-14}$ | $1.19\times10^{-22}$ | $1.41\times10^{-26}$ | $3.21\times10^{-06}$ |
| | | BP | $3.96\times10^{-18}$ | $3.10\times10^{-21}$ | $3.17\times10^{-20}$ | $1.39\times10^{-21}$ | $1.62\times10^{-24}$ | $2.99\times10^{-22}$ | $5.88\times10^{-22}$ | $1.49\times10^{-13}$ |
| | | CC | $7.76\times10^{-05}$ | $3.35\times10^{-14}$ | $2.98\times10^{-13}$ | $1.41\times10^{-11}$ | $1.52\times10^{-11}$ | $3.53\times10^{-01}$ | $2.32\times10^{-03}$ | $1.20\times10^{-01}$ |
| Rat | $Fmax$ | MF | $8.51\times10^{-18}$ | $3.25\times10^{-11}$ | $1.73\times10^{-11}$ | $4.62\times10^{-19}$ | $8.20\times10^{-01}$ | $6.32\times10^{-15}$ | $7.91\times10^{-16}$ | $4.98\times10^{-01}$ |
| | | BP | $5.04\times10^{-10}$ | $2.14\times10^{-11}$ | $4.67\times10^{-11}$ | $4.26\times10^{-14}$ | $6.72\times10^{-08}$ | $2.01\times10^{-06}$ | $3.11\times10^{-07}$ | $2.46\times10^{-01}$ |
| | | CC | $2.12\times10^{-08}$ | $3.22\times10^{-11}$ | $6.12\times10^{-12}$ | $2.22\times10^{-12}$ | $1.50\times10^{-08}$ | $1.23\times10^{-03}$ | $1.24\times10^{-02}$ | $8.58\times10^{-04}$ |
| | $AUPR$ | MF | $1.22\times10^{-22}$ | $6.41\times10^{-21}$ | $6.20\times10^{-20}$ | $1.43\times10^{-23}$ | $4.55\times10^{-09}$ | $2.09\times10^{-28}$ | $9.71\times10^{-27}$ | $2.84\times10^{-09}$ |
| | | BP | $1.03\times10^{-16}$ | $5.61\times10^{-20}$ | $3.12\times10^{-19}$ | $3.76\times10^{-20}$ | $1.05\times10^{-21}$ | $5.15\times10^{-14}$ | $5.67\times10^{-20}$ | $1.52\times10^{-03}$ |
| | | CC | $5.31\times10^{-09}$ | $2.13\times10^{-15}$ | $2.28\times10^{-15}$ | $6.98\times10^{-15}$ | $1.08\times10^{-14}$ | $5.16\times10^{-07}$ | $1.91\times10^{-02}$ | $5.95\times10^{-02}$ |
| Fly | $Fmax$ | MF | $1.60\times10^{-18}$ | $1.42\times10^{-15}$ | $1.19\times10^{-12}$ | $1.83\times10^{-20}$ | $6.27\times10^{-12}$ | $5.87\times10^{-12}$ | $5.21\times10^{-19}$ | $7.79\times10^{-06}$ |
| | | BP | $2.13\times10^{-12}$ | $5.73\times10^{-15}$ | $9.23\times10^{-13}$ | $2.09\times10^{-16}$ | $1.57\times10^{-14}$ | $6.39\times10^{-09}$ | $4.25\times10^{-16}$ | $4.74\times10^{-05}$ |
| | | CC | $1.00\times10^{-09}$ | $5.94\times10^{-15}$ | $4.35\times10^{-12}$ | $1.00\times10^{-14}$ | $3.24\times10^{-16}$ | $8.24\times10^{-03}$ | $8.18\times10^{-15}$ | $1.64\times10^{-05}$ |
| | $AUPR$ | MF | $3.30\times10^{-22}$ | $2.71\times10^{-22}$ | $3.56\times10^{-21}$ | $4.08\times10^{-24}$ | $6.16\times10^{-26}$ | $5.94\times10^{-25}$ | $5.07\times10^{-25}$ | $9.30\times10^{-11}$ |
| | | BP | $2.32\times10^{-15}$ | $1.09\times10^{-19}$ | $5.40\times10^{-18}$ | $1.61\times10^{-19}$ | $1.61\times10^{-25}$ | $2.45\times10^{-10}$ | $1.16\times10^{-19}$ | $3.01\times10^{-12}$ |
| | | CC | $2.59\times10^{-10}$ | $3.70\times10^{-19}$ | $1.35\times10^{-17}$ | $6.36\times10^{-18}$ | $9.80\times10^{-21}$ | $4.82\times10^{-02}$ | $1.16\times10^{-15}$ | $1.02\times10^{-04}$ |
| Budding Yeast | $Fmax$ | MF | $3.34\times10^{-16}$ | $4.96\times10^{-16}$ | $1.04\times10^{-11}$ | $4.28\times10^{-18}$ | $5.01\times10^{-15}$ | $5.42\times10^{-09}$ | $3.43\times10^{-20}$ | $8.27\times10^{-04}$ |
| | | BP | $8.17\times10^{-10}$ | $1.10\times10^{-14}$ | $1.31\times10^{-11}$ | $3.14\times10^{-14}$ | $1.76\times10^{-14}$ | $7.76\times10^{-01}$ | $1.91\times10^{-11}$ | $4.07\times10^{-02}$ |
| | | CC | $8.42\times10^{-02}$ | $2.97\times10^{-12}$ | $1.29\times10^{-07}$ | $1.34\times10^{-09}$ | $1.41\times10^{-08}$ | $9.25\times10^{-01}$ | $1.59\times10^{-02}$ | $8.00\times10^{-03}$ |
| | $AUPR$ | MF | $1.15\times10^{-17}$ | $3.63\times10^{-20}$ | $1.53\times10^{-18}$ | $4.30\times10^{-20}$ | $1.61\times10^{-22}$ | $1.23\times10^{-15}$ | $1.43\times10^{-25}$ | $5.33\times10^{-01}$ |
| | | BP | $6.37\times10^{-15}$ | $4.23\times10^{-21}$ | $4.02\times10^{-19}$ | $8.61\times10^{-20}$ | $2.08\times10^{-26}$ | $9.02\times10^{-11}$ | $1.77\times10^{-21}$ | $1.23\times10^{-09}$ |
| | | CC | $5.34\times10^{-17}$ | $3.81\times10^{-25}$ | $1.98\times10^{-23}$ | $2.54\times10^{-23}$ | $8.53\times10^{-22}$ | $1.22\times10^{-05}$ | $2.16\times10^{-23}$ | $4.61\times10^{-14}$ |
| Fission Yeast | $Fmax$ | MF | $5.83\times10^{-20}$ | $1.68\times10^{-16}$ | $8.72\times10^{-01}$ | $1.95\times10^{-20}$ | $6.87\times10^{-03}$ | $8.26\times10^{-13}$ | $1.14\times10^{-25}$ | $6.42\times10^{-01}$ |
| | | BP | $3.00\times10^{-12}$ | $2.56\times10^{-14}$ | $2.44\times10^{-10}$ | $3.25\times10^{-14}$ | $1.66\times10^{-07}$ | $2.30\times10^{-04}$ | $3.11\times10^{-18}$ | $1.88\times10^{-02}$ |
| | | CC | $5.15\times10^{-13}$ | $1.71\times10^{-16}$ | $3.83\times10^{-13}$ | $3.41\times10^{-15}$ | $3.03\times10^{-12}$ | $4.83\times10^{-05}$ | $7.40\times10^{-16}$ | $3.99\times10^{-11}$ |
| | $AUPR$ | MF | $6.64\times10^{-20}$ | $4.96\times10^{-20}$ | $4.11\times10^{-18}$ | $1.21\times10^{-20}$ | $4.03\times10^{-09}$ | $1.86\times10^{-16}$ | $2.11\times10^{-28}$ | $1.63\times10^{-07}$ |
| | | BP | $1.93\times10^{-18}$ | $3.65\times10^{-22}$ | $2.38\times10^{-19}$ | $5.18\times10^{-21}$ | $7.43\times10^{-17}$ | $1.56\times10^{-11}$ | $3.91\times10^{-27}$ | $1.41\times10^{-08}$ |
| | | CC | $5.39\times10^{-14}$ | $1.01\times10^{-19}$ | $9.20\times10^{-18}$ | $2.48\times10^{-18}$ | $9.57\times10^{-22}$ | $3.94\times10^{-01}$ | $1.37\times10^{-18}$ | $5.73\times10^{-07}$ |
| Nematoda | $Fmax$ | MF | $3.62\times10^{-14}$ | $6.79\times10^{-11}$ | $8.72\times10^{-01}$ | $8.85\times10^{-16}$ | $8.91\times10^{-01}$ | $7.73\times10^{-08}$ | $6.60\times10^{-18}$ | $7.18\times10^{-03}$ |
| | | BP | $2.31\times10^{-08}$ | $9.59\times10^{-12}$ | $9.80\times10^{-10}$ | $1.39\times10^{-12}$ | $4.25\times10^{-12}$ | $5.61\times10^{-03}$ | $1.40\times10^{-09}$ | $7.70\times10^{-01}$ |
| | | CC | $8.47\times10^{-08}$ | $6.57\times10^{-15}$ | $4.39\times10^{-12}$ | $9.90\times10^{-13}$ | $3.53\times10^{-07}$ | $1.69\times10^{-02}$ | $1.04\times10^{-10}$ | $1.90\times10^{-03}$ |
| | $AUPR$ | MF | $2.34\times10^{-17}$ | $1.28\times10^{-17}$ | $2.23\times10^{-13}$ | $1.34\times10^{-18}$ | $1.00\times10^{-01}$ | $2.24\times10^{-01}$ | $3.21\times10^{-23}$ | $5.07\times10^{-02}$ |
| | | BP | $6.74\times10^{-14}$ | $5.92\times10^{-19}$ | $4.85\times10^{-17}$ | $3.38\times10^{-18}$ | $3.54\times10^{-22}$ | $6.00\times10^{-08}$ | $1.48\times10^{-17}$ | $5.85\times10^{-02}$ |
| | | CC | $1.33\times10^{-09}$ | $1.28\times10^{-18}$ | $3.85\times10^{-16}$ | $3.71\times10^{-16}$ | $2.44\times10^{-15}$ | $5.88\times10^{-01}$ | $1.81\times10^{-10}$ | $9.70\times10^{-02}$ |

**Table S6 The p-values between EGN and other six GO prediction methods for $Fmax$ and $AUPR$ on 98 non-coding genes**

| Measure | GO aspect | (EGN, EPGP) | (EGN, GSAGP) | (EGN, NGP) | (EGN, GN) | (EGN, EN) | (EGN, EG) |
|---|---|---|---|---|---|---|---|
| $WAFmax$ | MF | $1.09\times10^{-12}$ | $4.01\times10^{-17}$ | $5.48\times10^{-18}$ | $4.01\times10^{-17}$ | $1.95\times10^{-12}$ | $4.16\times10^{-10}$ |
| | BP | $3.86\times10^{-10}$ | $3.36\times10^{-18}$ | $1.07\times10^{-16}$ | $7.98\times10^{-17}$ | $3.86\times10^{-10}$ | $1.39\times10^{-03}$ |
| | CC | $1.73\times10^{-05}$ | $1.35\times10^{-13}$ | $1.35\times10^{-13}$ | $1.17\times10^{-11}$ | $1.80\times10^{-03}$ | $2.80\times10^{-05}$ |
| $WAAUPR$ | MF | $4.83\times10^{-20}$ | $3.39\times10^{-25}$ | $2.87\times10^{-25}$ | $4.30\times10^{-24}$ | $1.13\times10^{-19}$ | $6.30\times10^{-17}$ |
| | BP | $5.90\times10^{-09}$ | $1.51\times10^{-23}$ | $1.99\times10^{-21}$ | $9.56\times10^{-21}$ | $1.38\times10^{-09}$ | $5.88\times10^{-11}$ |
| | CC | $1.56\times10^{-11}$ | $8.53\times10^{-25}$ | $2.93\times10^{-24}$ | $3.07\times10^{-22}$ | $1.56\times10^{-11}$ | $1.01\times10^{-05}$ |

**Table S7 The details of training and test dataset for 7 species in CAFA3 dataset**

| Species | NTR[1] | NTE[1] | NGT_MF[2] | NGT_BP[3] | NGT_CC[4] |
|---|---|---|---|---|---|
| Human | 13639 | 1020 | 3611 | 10680 | 1348 |
| Mouse | 9220 | 277 | 2374 | 10949 | 1004 |
| Arabidopsis | 7550 | 470 | 1958 | 4230 | 516 |
| Rat | 3645 | 75 | 2198 | 6684 | 784 |
| Fly | 2420 | 196 | 1263 | 4875 | 693 |
| Budding Yeast | 4061 | 23 | 1986 | 4511 | 877 |
| Fission Yeast | 4237 | 372 | 1237 | 3824 | 626 |

[1]NTR/NTE: the number of proteins in training/test datasets

[2]NGT_MF: the total number of MF terms in training and test datasets

[3]NGT_BP: the total number of BP terms in training and test datasets

[4]NGT_CC: the total number of CC terms in training and test datasets

**Table S8 The p-values between TNP and other five expression profile-based methods for $Fmax$ and $AUPR$ on 2433 proteins of 7 species from CAFA3 test dataset**

| Measure | GO aspect | (TNP, MR) | (TNP, PCC) | (TNP, MLC) | (TNP, SRC) | (TNP, ED) |
|---|---|---|---|---|---|---|
| | MF | $8.07\times10^{-09}$ | $4.92\times10^{-09}$ | $2.25\times10^{-15}$ | $7.16\times10^{-09}$ | $1.29\times10^{-09}$ |
| $Fmax$ | BP | $2.87\times10^{-07}$ | $3.95\times10^{-09}$ | $5.38\times10^{-14}$ | $2.07\times10^{-08}$ | $8.28\times10^{-09}$ |
| | CC | $2.14\times10^{-07}$ | $8.38\times10^{-09}$ | $3.53\times10^{-12}$ | $1.45\times10^{-07}$ | $3.72\times10^{-09}$ |
| | MF | $1.41\times10^{-01}$ | $3.10\times10^{-05}$ | $2.41\times10^{-11}$ | $3.13\times10^{-04}$ | $1.15\times10^{-05}$ |
| $AUPR$ | BP | $2.49\times10^{-05}$ | $4.09\times10^{-07}$ | $2.76\times10^{-13}$ | $6.15\times10^{-06}$ | $1.15\times10^{-06}$ |
| | CC | $8.01\times10^{-01}$ | $3.59\times10^{-08}$ | $7.11\times10^{-05}$ | $4.46\times10^{-02}$ | $1.04\times10^{-08}$ |

**Table S9 The p-values between TNP and other five expression profile-based methods for $Fmax$ and $AUPR$ on CAFA3 test dataset for each of 7 species**

| Species | Measure | GO aspect | (TNP, MR) | (TNP, PCC) | (TNP, MLC) | (TNP, SRC) | (TNP, ED) |
|---|---|---|---|---|---|---|---|
| Human | $Fmax$ | MF | $1.69\times10^{-02}$ | $4.86\times10^{-05}$ | $2.09\times10^{-04}$ | $4.09\times10^{-03}$ | $9.32\times10^{-06}$ |
| | | BP | $1.48\times10^{-02}$ | $4.96\times10^{-06}$ | $2.54\times10^{-09}$ | $1.57\times10^{-03}$ | $2.15\times10^{-05}$ |
| | | CC | $3.58\times10^{-04}$ | $6.87\times10^{-07}$ | $5.46\times10^{-09}$ | $1.09\times10^{-05}$ | $2.79\times10^{-07}$ |
| | $AUPR$ | MF | $4.64\times10^{-04}$ | $2.80\times10^{-04}$ | $9.63\times10^{-01}$ | $4.54\times10^{-05}$ | $3.01\times10^{-05}$ |
| | | BP | $7.63\times10^{-02}$ | $3.40\times10^{-04}$ | $3.79\times10^{-08}$ | $1.37\times10^{-01}$ | $3.87\times10^{-03}$ |
| | | CC | $9.28\times10^{-02}$ | $4.54\times10^{-08}$ | $4.59\times10^{-03}$ | $9.28\times10^{-04}$ | $4.54\times10^{-08}$ |
| Mouse | $Fmax$ | MF | $6.45\times10^{-06}$ | $3.51\times10^{-08}$ | $8.42\times10^{-09}$ | $3.32\times10^{-07}$ | $1.70\times10^{-07}$ |
| | | BP | $8.47\times10^{-08}$ | $3.37\times10^{-08}$ | $1.05\times10^{-11}$ | $2.31\times10^{-07}$ | $1.39\times10^{-07}$ |
| | | CC | $1.98\times10^{-04}$ | $3.74\times10^{-03}$ | $2.39\times10^{-04}$ | $5.25\times10^{-03}$ | $1.23\times10^{-04}$ |
| | $AUPR$ | MF | $7.05\times10^{-01}$ | $9.43\times10^{-07}$ | $3.90\times10^{-06}$ | $8.26\times10^{-02}$ | $2.30\times10^{-06}$ |
| | | BP | $3.54\times10^{-02}$ | $1.55\times10^{-05}$ | $8.05\times10^{-08}$ | $2.27\times10^{-01}$ | $2.01\times10^{-06}$ |
| | | CC | $8.23\times10^{-09}$ | $4.69\times10^{-07}$ | $2.96\times10^{-10}$ | $2.28\times10^{-08}$ | $8.67\times10^{-07}$ |
| Arabidopsis | $Fmax$ | MF | $6.05\times10^{-06}$ | $2.64\times10^{-06}$ | $6.28\times10^{-08}$ | $1.56\times10^{-04}$ | $7.72\times10^{-07}$ |
| | | BP | $1.87\times10^{-02}$ | $6.95\times10^{-05}$ | $6.53\times10^{-12}$ | $1.41\times10^{-05}$ | $1.10\times10^{-04}$ |
| | | CC | $7.25\times10^{-05}$ | $6.74\times10^{-07}$ | $2.14\times10^{-05}$ | $6.90\times10^{-06}$ | $1.53\times10^{-07}$ |
| | $AUPR$ | MF | $2.48\times10^{-03}$ | $2.48\times10^{-03}$ | $3.16\times10^{-06}$ | $5.81\times10^{-01}$ | $3.41\times10^{-04}$ |
| | | BP | $3.06\times10^{-03}$ | $1.24\times10^{-05}$ | $1.32\times10^{-13}$ | $4.61\times10^{-08}$ | $3.89\times10^{-05}$ |
| | | CC | $6.33\times10^{-03}$ | $1.60\times10^{-04}$ | $4.47\times10^{-06}$ | $3.20\times10^{-03}$ | $2.76\times10^{-04}$ |
| Rat | $Fmax$ | MF | $5.23\times10^{-01}$ | $3.58\times10^{-01}$ | $5.07\times10^{-04}$ | $9.21\times10^{-03}$ | $2.41\times10^{-01}$ |
| | | BP | $3.08\times10^{-01}$ | $8.43\times10^{-06}$ | $1.94\times10^{-02}$ | $1.56\times10^{-01}$ | $4.71\times10^{-06}$ |
| | | CC | $2.76\times10^{-02}$ | $4.50\times10^{-02}$ | $3.07\times10^{-04}$ | $6.25\times10^{-01}$ | $1.93\times10^{-02}$ |
| | $AUPR$ | MF | $4.14\times10^{-01}$ | $4.14\times10^{-01}$ | $3.48\times10^{-03}$ | $9.47\times10^{-02}$ | $1.37\times10^{-01}$ |
| | | BP | $1.12\times10^{-04}$ | $2.88\times10^{-02}$ | $5.56\times10^{-04}$ | $1.32\times10^{-06}$ | $1.90\times10^{-03}$ |
| | | CC | $6.57\times10^{-03}$ | $1.87\times10^{-02}$ | $2.39\times10^{-03}$ | $9.17\times10^{-02}$ | $6.57\times10^{-03}$ |
| Fly | $Fmax$ | MF | $6.79\times10^{-03}$ | $1.08\times10^{-01}$ | $5.13\times10^{-05}$ | $8.70\times10^{-04}$ | $1.23\times10^{-02}$ |
| | | BP | $3.62\times10^{-03}$ | $5.03\times10^{-02}$ | $3.94\times10^{-07}$ | $8.67\times10^{-04}$ | $2.66\times10^{-03}$ |
| | | CC | $2.40\times10^{-04}$ | $1.48\times10^{-04}$ | $1.36\times10^{-06}$ | $2.42\times10^{-05}$ | $4.67\times10^{-06}$ |
| | $AUPR$ | MF | $5.84\times10^{-06}$ | $1.03\times10^{-05}$ | $8.06\times10^{-11}$ | $1.48\times10^{-08}$ | $2.05\times10^{-06}$ |
| | | BP | $1.16\times10^{-03}$ | $2.01\times10^{-04}$ | $5.38\times10^{-10}$ | $1.21\times10^{-08}$ | $8.70\times10^{-03}$ |
| | | CC | $3.51\times10^{-04}$ | $6.82\times10^{-06}$ | $1.02\times10^{-06}$ | $1.20\times10^{-05}$ | $6.14\times10^{-07}$ |
| Budding Yeast | $Fmax$ | MF | $2.63\times10^{-04}$ | $8.95\times10^{-02}$ | $5.03\times10^{-04}$ | $3.36\times10^{-06}$ | $1.90\times10^{-03}$ |
| | | BP | $1.02\times10^{-01}$ | $1.28\times10^{-06}$ | $1.28\times10^{-02}$ | $2.08\times10^{-05}$ | $2.99\times10^{-06}$ |
| | | CC | $2.39\times10^{-03}$ | $1.17\times10^{-02}$ | $2.89\times10^{-04}$ | $1.67\times10^{-03}$ | $1.17\times10^{-02}$ |
| | $AUPR$ | MF | $8.79\times10^{-01}$ | $4.29\times10^{-02}$ | $9.93\times10^{-06}$ | $4.20\times10^{-08}$ | $4.29\times10^{-02}$ |
| | | BP | $6.74\times10^{-01}$ | $6.74\times10^{-01}$ | $6.41\times10^{-01}$ | $4.03\times10^{-04}$ | $7.47\times10^{-01}$ |
| | | CC | $2.55\times10^{-05}$ | $9.48\times10^{-03}$ | $6.52\times10^{-01}$ | $1.17\times10^{-05}$ | $2.74\times10^{-03}$ |
| Fission Yeast | $Fmax$ | MF | $7.90\times10^{-01}$ | $5.46\times10^{-01}$ | $6.95\times10^{-03}$ | $3.88\times10^{-01}$ | $7.58\times10^{-01}$ |
| | | BP | $2.72\times10^{-10}$ | $7.08\times10^{-11}$ | $2.30\times10^{-16}$ | $3.86\times10^{-11}$ | $7.08\times10^{-11}$ |
| | | CC | $3.42\times10^{-02}$ | $4.53\times10^{-05}$ | $4.31\times10^{-05}$ | $1.38\times10^{-04}$ | $4.53\times10^{-05}$ |
| | $AUPR$ | MF | $9.89\times10^{-06}$ | $3.37\times10^{-06}$ | $7.07\times10^{-09}$ | $3.31\times10^{-05}$ | $3.37\times10^{-06}$ |
| | | BP | $7.06\times10^{-13}$ | $3.08\times10^{-13}$ | $1.59\times10^{-19}$ | $2.49\times10^{-13}$ | $3.08\times10^{-13}$ |
| | | CC | $3.37\times10^{-04}$ | $9.72\times10^{-03}$ | $6.07\times10^{-03}$ | $4.84\times10^{-03}$ | $8.15\times10^{-03}$ |

**Table S10 The p-values between TripletGO and other six GO prediction methods for $Fmax$ and $AUPR$ on 2,433 proteins of 7 species from CAFA3 test dataset**

| Measure | GO aspect | (TripletGO, EPGP) | (TripletGO, GSAGP) | (TripletGO, PSAGP) | (TripletGO, NGP) | (TripletGO, DeepGO) | (TripletGO, FunFams) |
|---|---|---|---|---|---|---|---|
| $Fmax$ | MF | $4.75\times10^{-19}$ | $4.07\times10^{-16}$ | $9.05\times10^{-02}$ | $8.09\times10^{-20}$ | $8.74\times10^{-18}$ | $2.07\times10^{-07}$ |
| | BP | $2.79\times10^{-08}$ | $1.42\times10^{-11}$ | $5.90\times10^{-07}$ | $2.12\times10^{-13}$ | $5.44\times10^{-09}$ | $1.83\times10^{-07}$ |
| | CC | $5.99\times10^{-10}$ | $1.56\times10^{-14}$ | $9.02\times10^{-10}$ | $1.31\times10^{-12}$ | $1.63\times10^{-10}$ | $7.58\times10^{-14}$ |
| $AUPR$ | MF | $5.58\times10^{-22}$ | $1.22\times10^{-21}$ | $3.60\times10^{-19}$ | $1.36\times10^{-22}$ | $1.88\times10^{-20}$ | $1.69\times10^{-18}$ |
| | BP | $2.12\times10^{-15}$ | $1.12\times10^{-18}$ | $1.93\times10^{-14}$ | $2.21\times10^{-18}$ | $4.69\times10^{-16}$ | $1.37\times10^{-17}$ |
| | CC | $2.09\times10^{-11}$ | $2.11\times10^{-19}$ | $5.95\times10^{-17}$ | $2.38\times10^{-16}$ | $1.46\times10^{-11}$ | $1.62\times10^{-18}$ |

**Table S11 The numbers of genes with GO annotation of three aspects for 20 species**

| Database | Species | Version | Gene number | Sample number | GO number | MF number | BP number | CC number |
|---|---|---|---|---|---|---|---|---|
| COXPRESdb | Nematoda | Cel-m.c4-0 | 17256 | 1780 | 3154 | 1254 | 2705 | 2018 |
| | Dog | Cfa-m.c3-0 | 16214 | 777 | 96 | 31 | 54 | 79 |
| | Fly | Dme-m.c4-0 | 12626 | 4209 | 5317 | 2729 | 4874 | 3495 |
| | Zebrafish | Dre-m.c4-0 | 10112 | 1423 | 2477 | 541 | 2324 | 414 |
| | Chicken | Gga-m.c4-0 | 13757 | 1502 | 502 | 215 | 383 | 337 |
| | Human | Hsa-m2.c3-0 | 20199 | 27655 | 14706 | 9281 | 12362 | 13278 |
| | Monkey | Mcc-m.c3-0 | 15782 | 1006 | 0 | 0 | 0 | 0 |
| | Mouse | Mmu-m.c4-0 | 20962 | 42916 | 10564 | 5646 | 8909 | 7621 |
| | Rat | Rno-m.c4-0 | 13751 | 42752 | 5409 | 3594 | 4387 | 4135 |
| | Budding yeast | Sce-m.c3-0 | 4461 | 3593 | 4107 | 3130 | 3934 | 3402 |
| | Fission yeast | Spo-m.c3-0 | 4881 | 166 | 2743 | 1303 | 2339 | 1877 |
| ATTED-II | Arabidopsis | Ath-m.c8-0 | 20819 | 12686 | 11602 | 5090 | 7927 | 8656 |
| | Field mustard | Bra-r.c3-0 | 26339 | 164 | 0 | 0 | 0 | 0 |
| | Soybean | Gma-m.c4-0 | 15746 | 1022 | 0 | 0 | 0 | 0 |
| | Medicago | Mtr-m.c4-1 | 20376 | 780 | 0 | 0 | 0 | 0 |
| | Rice | Osa-m.c7-0 | 19867 | 1775 | 82 | 59 | 69 | 55 |
| | Poplar | Ppo-m.c3-0 | 21910 | 557 | 0 | 0 | 0 | 0 |
| | Tomato | Sly-m.c4-0 | 5721 | 392 | 0 | 0 | 0 | 0 |
| | Grape | Vvi-m.c4-0 | 9421 | 258 | 0 | 0 | 0 | 0 |
| | Maize | Zma-m.c4-0 | 10777 | 606 | 0 | 0 | 0 | 0 |

Gene number: the total number of genes in a species.
Sample number: the number of experimental samples in microarray technology.
GO number: the number of genes with GO annotation in a species.
MF/BP/CC number: the number of genes with MF/BP/CC GO annotation in a species.

**Table S12 The details of 8 benchmark datasets constructed in our work**

| Species | NTR[1] | NEV[1] | NTE[1] | NGT_MF[2] | NGT_BP[3] | NGT_CC[4] |
|---|---|---|---|---|---|---|
| Human | 12501 | 735 | 1470 | 3841 | 11674 | 1505 |
| Mouse | 8965 | 527 | 1054 | 2735 | 12035 | 1188 |
| Arabidopsis | 9862 | 580 | 1160 | 2245 | 4787 | 563 |
| Rat | 4599 | 270 | 540 | 2542 | 7917 | 954 |
| Fly | 4521 | 265 | 531 | 1783 | 6123 | 857 |
| Budding Yeast | 3492 | 205 | 410 | 2025 | 4525 | 899 |
| Fission Yeast | 2332 | 137 | 274 | 1426 | 3885 | 720 |
| Nematoda | 2682 | 157 | 315 | 1160 | 4042 | 539 |

[1]NTR/NEV/NTE: the number of genes in training/validation/test datasets

[2]NGT_MF: the total number of MF terms in training, validation and test datasets

[3]NGT_BP: the total number of BP terms in training, validation and test datasets

[4]NGT_CC: the total number of CC terms in training, validation and test datasets

**Table S13 The values of $\alpha$, $h$, $margin$, and $c_f$ on the benchmark datasets for 8 species**

| Species | GO aspect | $\alpha$ | $h$ | $margin$ | $c_f$ |
|---|---|---|---|---|---|
| Human | MF | 5 | 1000 | 0.01 | 0.90 |
| | BP | 5 | 1000 | 0.01 | 0.90 |
| | CC | 10 | 1000 | 0.01 | 0.95 |
| Mouse | MF | 5 | 1000 | 0.01 | 0.90 |
| | BP | 3 | 1000 | 0.01 | 0.90 |
| | CC | 2 | 1000 | 0.01 | 0.95 |
| Arabidopsis | MF | 5 | 1000 | 0.01 | 0.90 |
| | BP | 5 | 1000 | 0.01 | 0.90 |
| | CC | 5 | 1000 | 0.01 | 0.95 |
| Rat | MF | 3 | 1000 | 0.01 | 0.90 |
| | BP | 5 | 1000 | 0.01 | 0.90 |
| | CC | 5 | 1000 | 0.01 | 0.95 |
| Fly | MF | 3 | 1000 | 0.01 | 0.90 |
| | BP | 5 | 1000 | 0.01 | 0.90 |
| | CC | 5 | 1000 | 0.01 | 0.95 |
| Budding Yeast | MF | 3 | 1000 | 0.01 | 0.90 |
| | BP | 5 | 1000 | 0.01 | 0.90 |
| | CC | 5 | 1000 | 0.01 | 0.95 |
| Fission Yeast | MF | 3 | - | 0.01 | 0.90 |
| | BP | 5 | - | 0.01 | 0.90 |
| | CC | 5 | - | 0.01 | 0.95 |
| Nematoda | MF | 5 | 1000 | 0.01 | 0.90 |
| | BP | 5 | 1000 | 0.01 | 0.90 |
| | CC | 5 | 1000 | 0.01 | 0.95 |

‘-’ means that the PCA is not executed in the corresponding species

### Supporting Figures

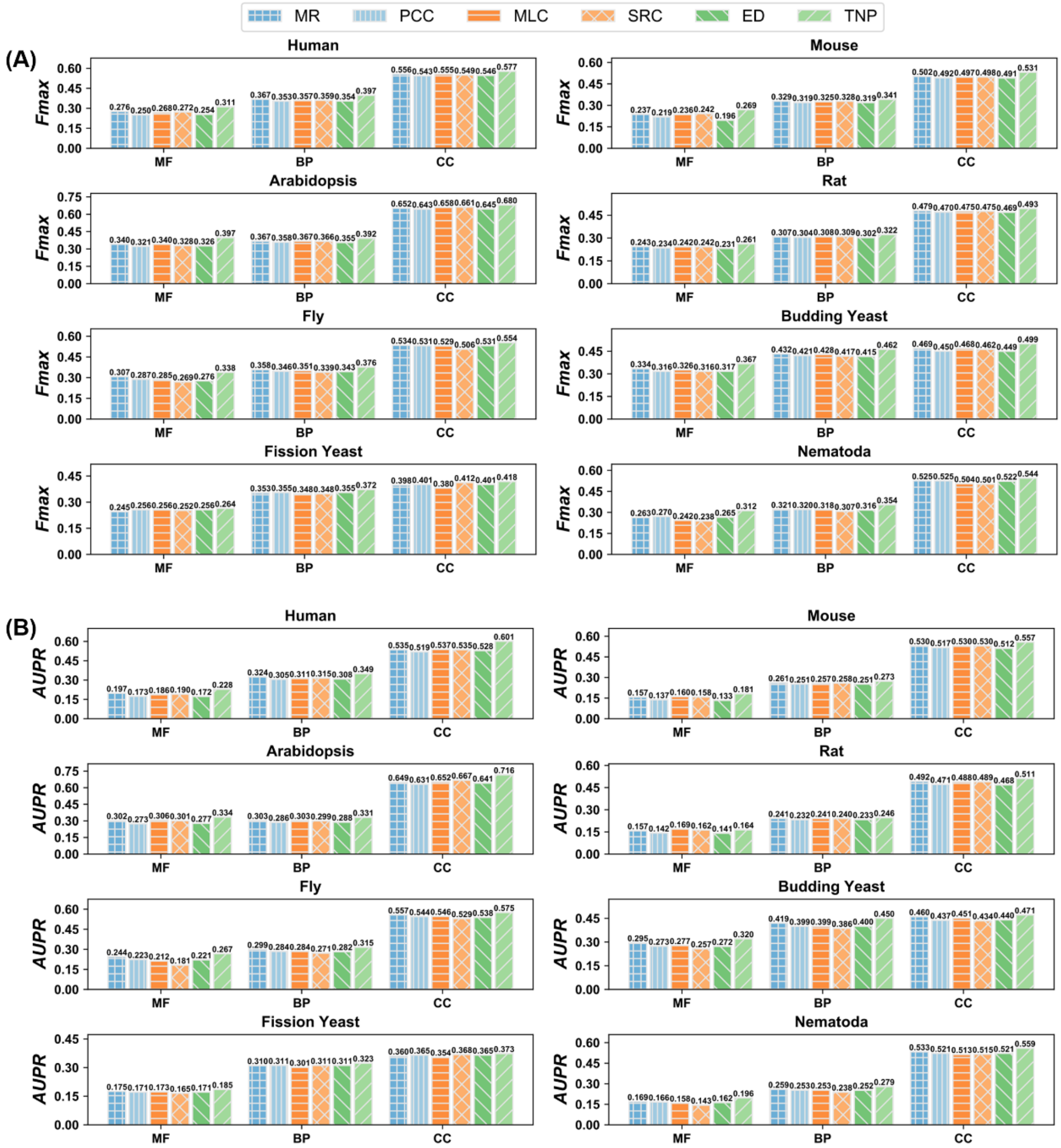


**Figure S1 The performance of six expression profile-based methods on the test datasets for 8 species**

**A**. The *Fmax* values of six methods for 8 species. **B**. The *AUPR* values of six methods for 8 species.

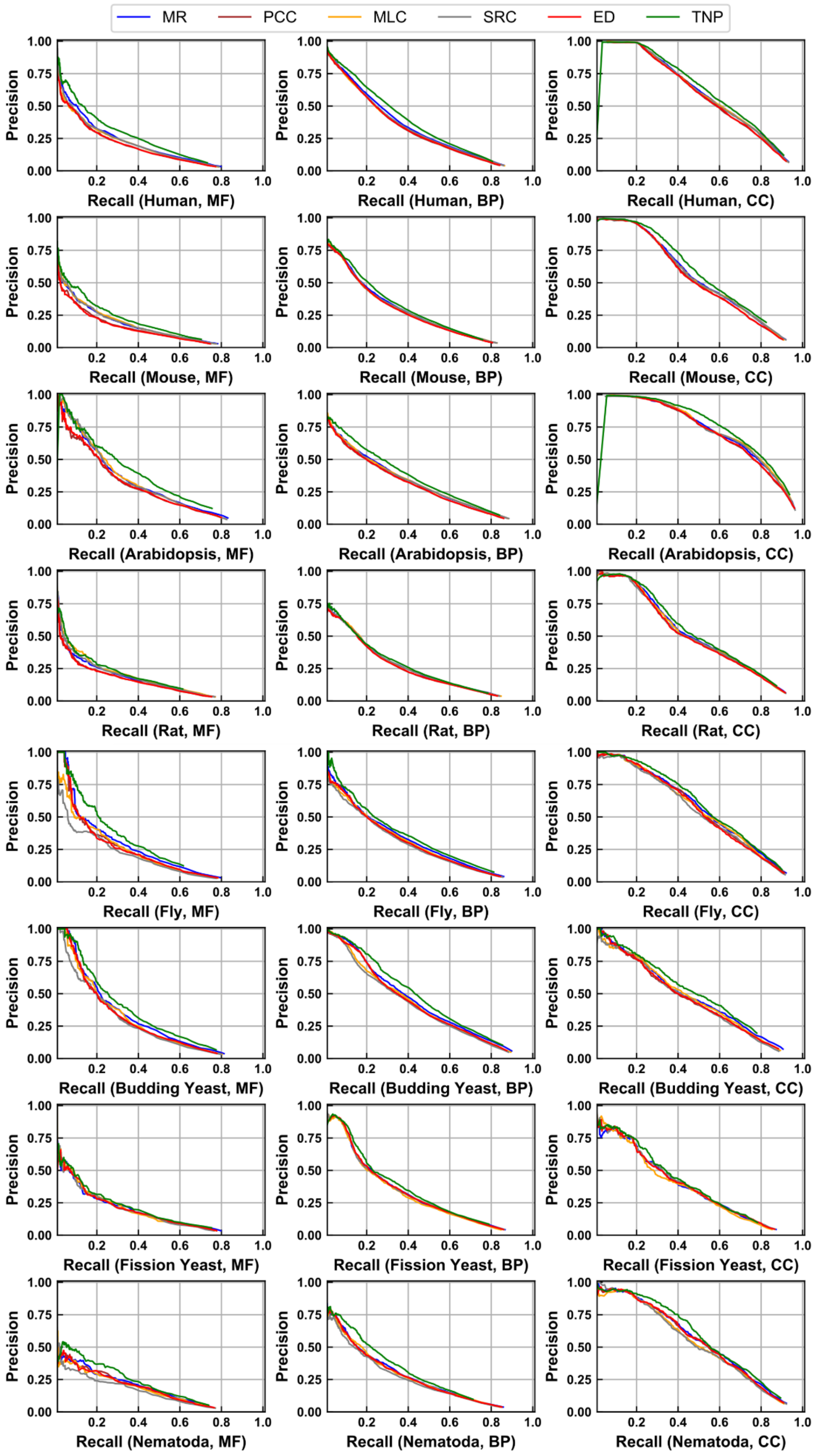


**Figure S2 The precision-recall curves of six expression profile-based methods on the test datasets for 8 species**

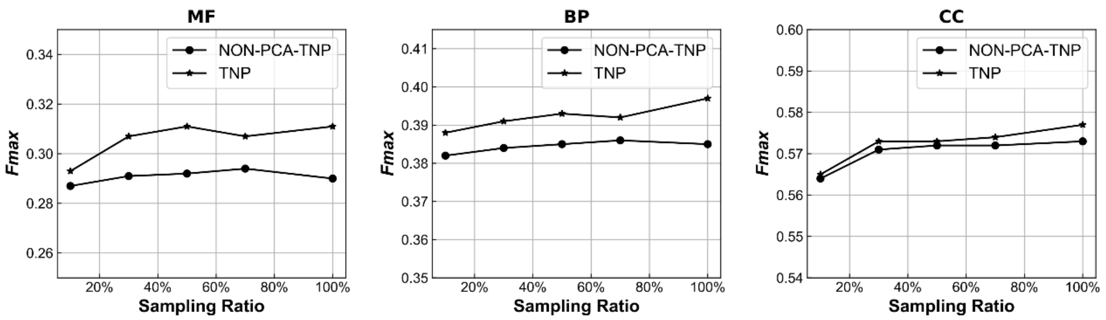


**Figure S3 Variation curves of $Fmax$ values of TNP and NON-PCA-TNP on the test dataset of human species versus the sampling ratios in expression data**

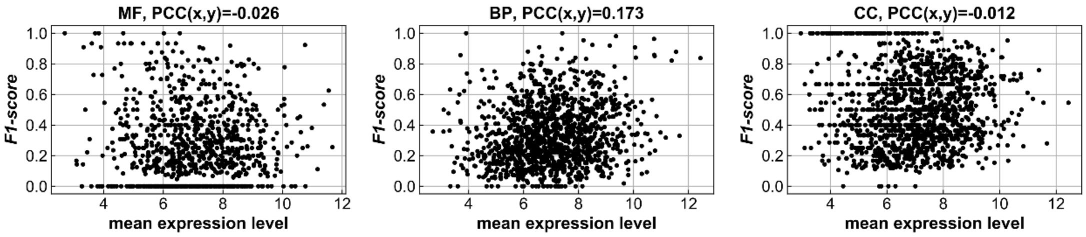


**Figure S4 The scattering plots of mean expression level versus F1-scores for 1470 human test genes by TNP**

PCC (x, y) means the PCC value between mean expression level and F1-score for all test genes

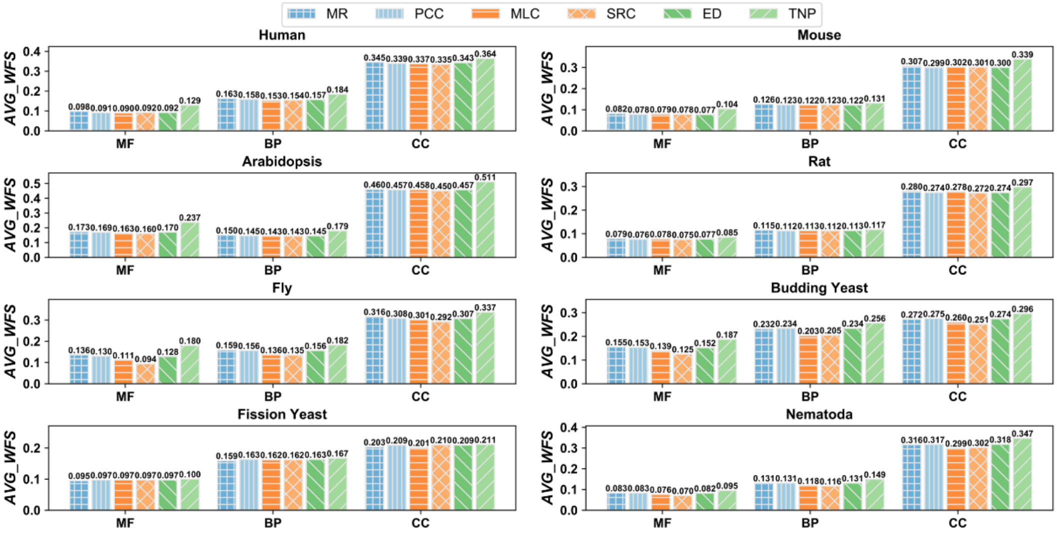


**Figure S5 The $AVG\_WFS$ values of six measures for three GO aspects in 8 individual species**

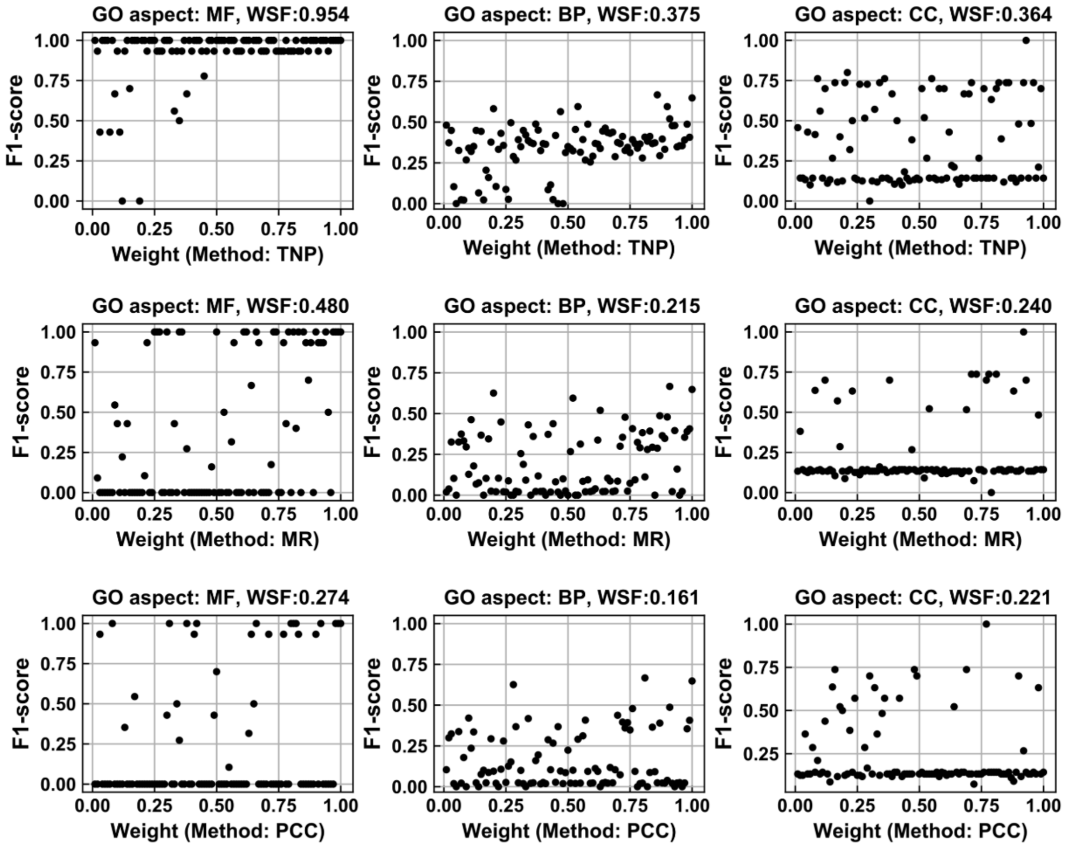


**Figure S6 The scattering plots of weights versus F1-scores of 100 templates for the gene MIRLET7C over TNP, MR, and PCC**

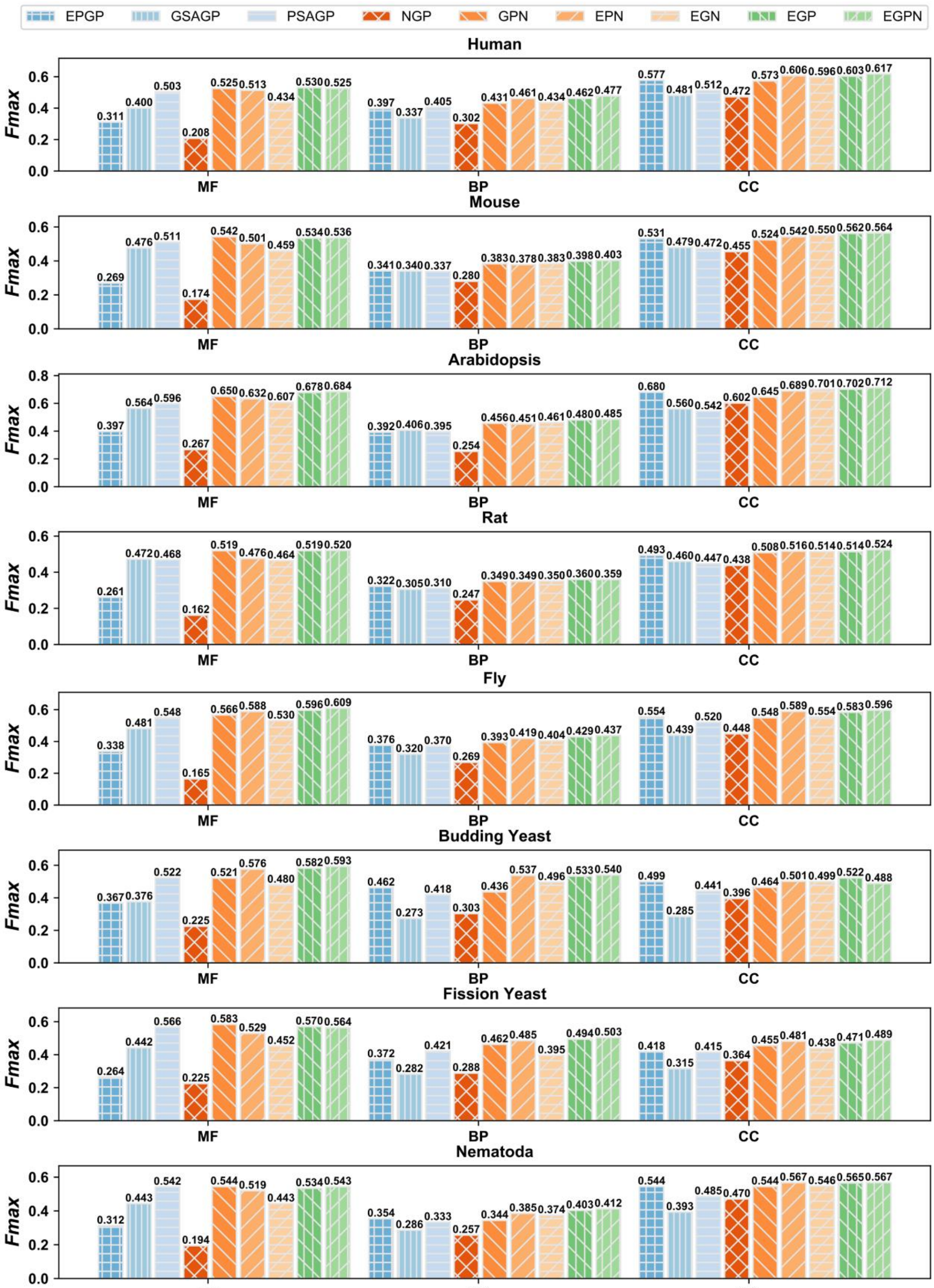


**Figure S7 The *Fmax* values of nine GO prediction methods on the test datasets for 8 species**

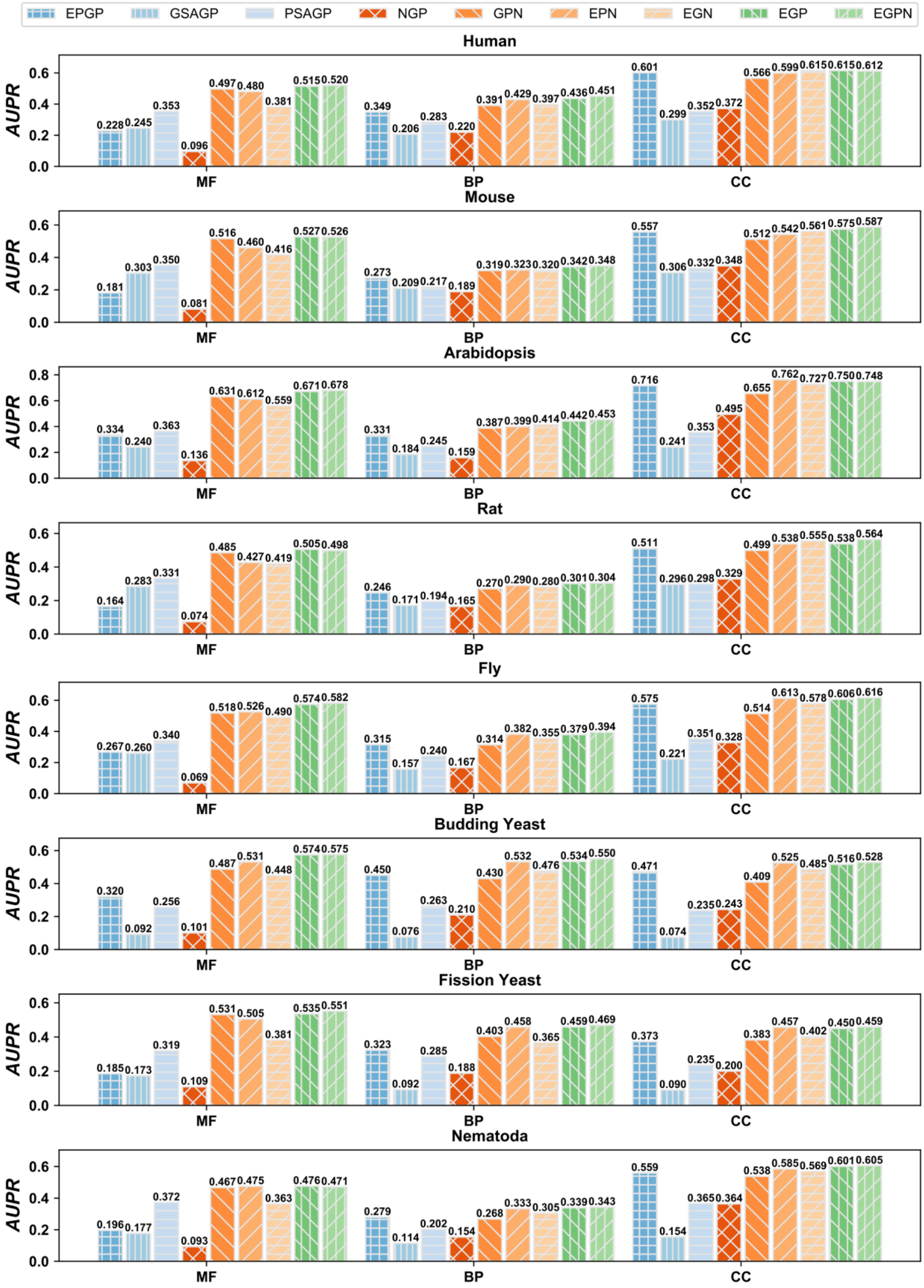


**Figure S8 The *AUPR* values of nine GO prediction methods on the test datasets for 8 species**

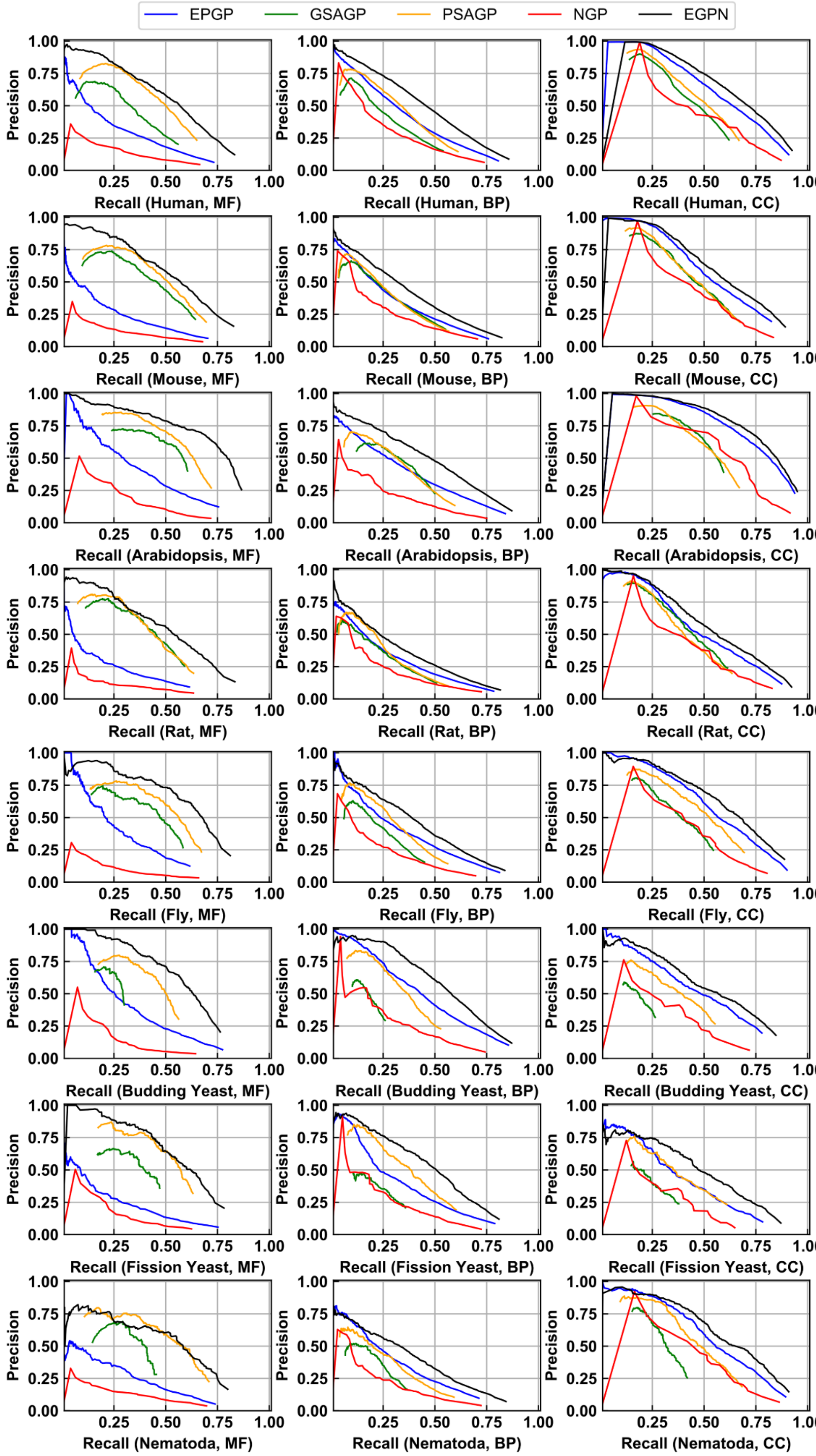


**Figure S9 The precision-recall curves of five GO prediction methods on the test datasets for 8 species**

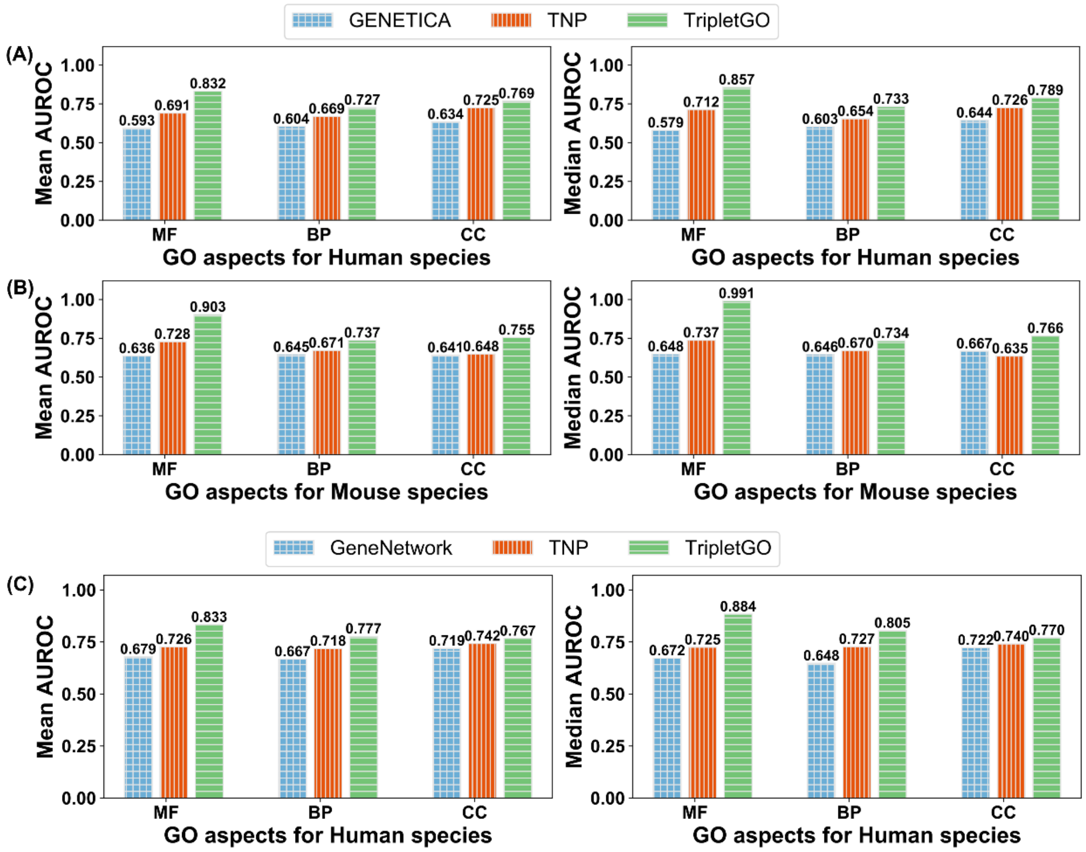


**Figure S10 Comparison of mean and median AUROC values of three GO aspects by different methods on the common dataset**

**A**. GENETICA, TNP and TripletGO on human; **B**. GENETICA, TNP and TripletGO on mouse; **C**. GeneNetwork, TNP and TripletGO on human.

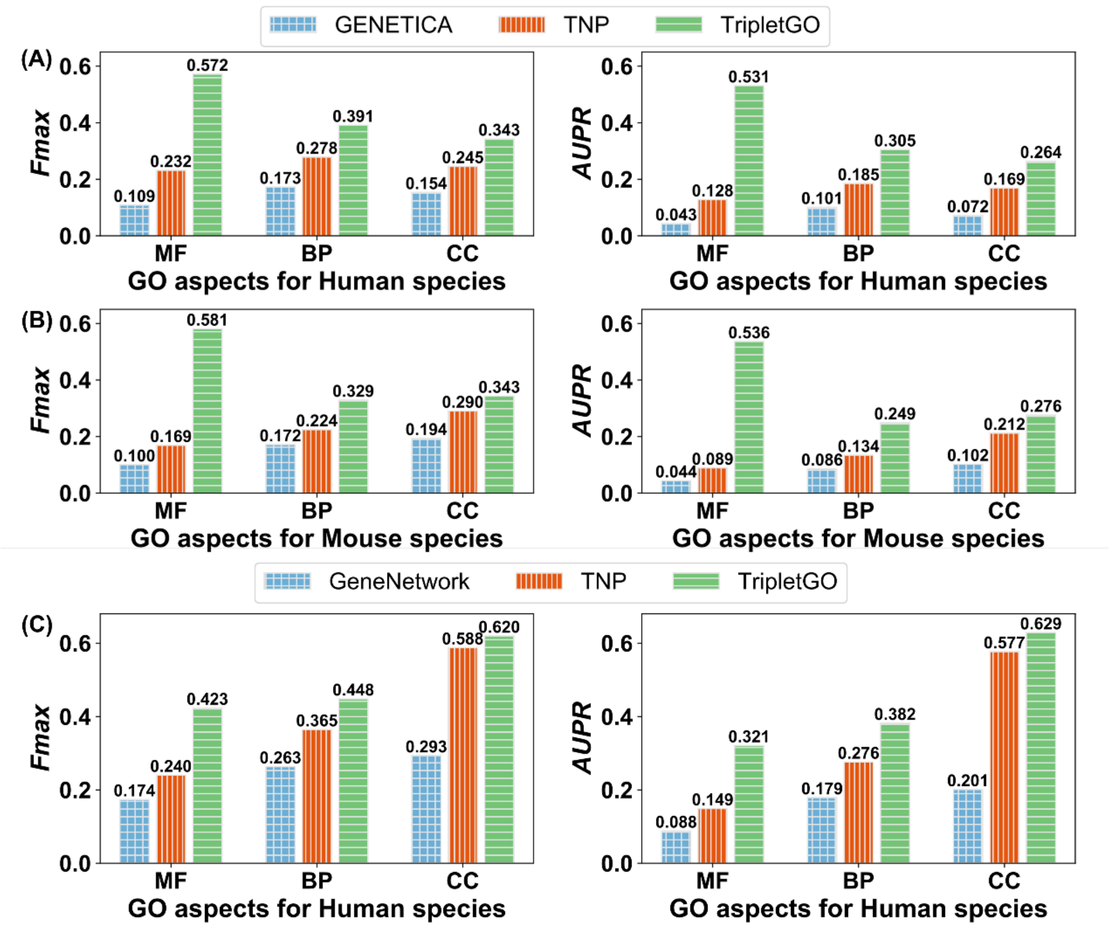


**Figure S11 Comparison of $Fmax$ and $AUPR$ values of three GO aspects by different methods on the common dataset**

**A**. GENETICA, TNP and TripletGO on human; **B**. GENETICA, TNP and TripletGO on mouse; **C**. GeneNetwork, TNP and TripletGO on human.

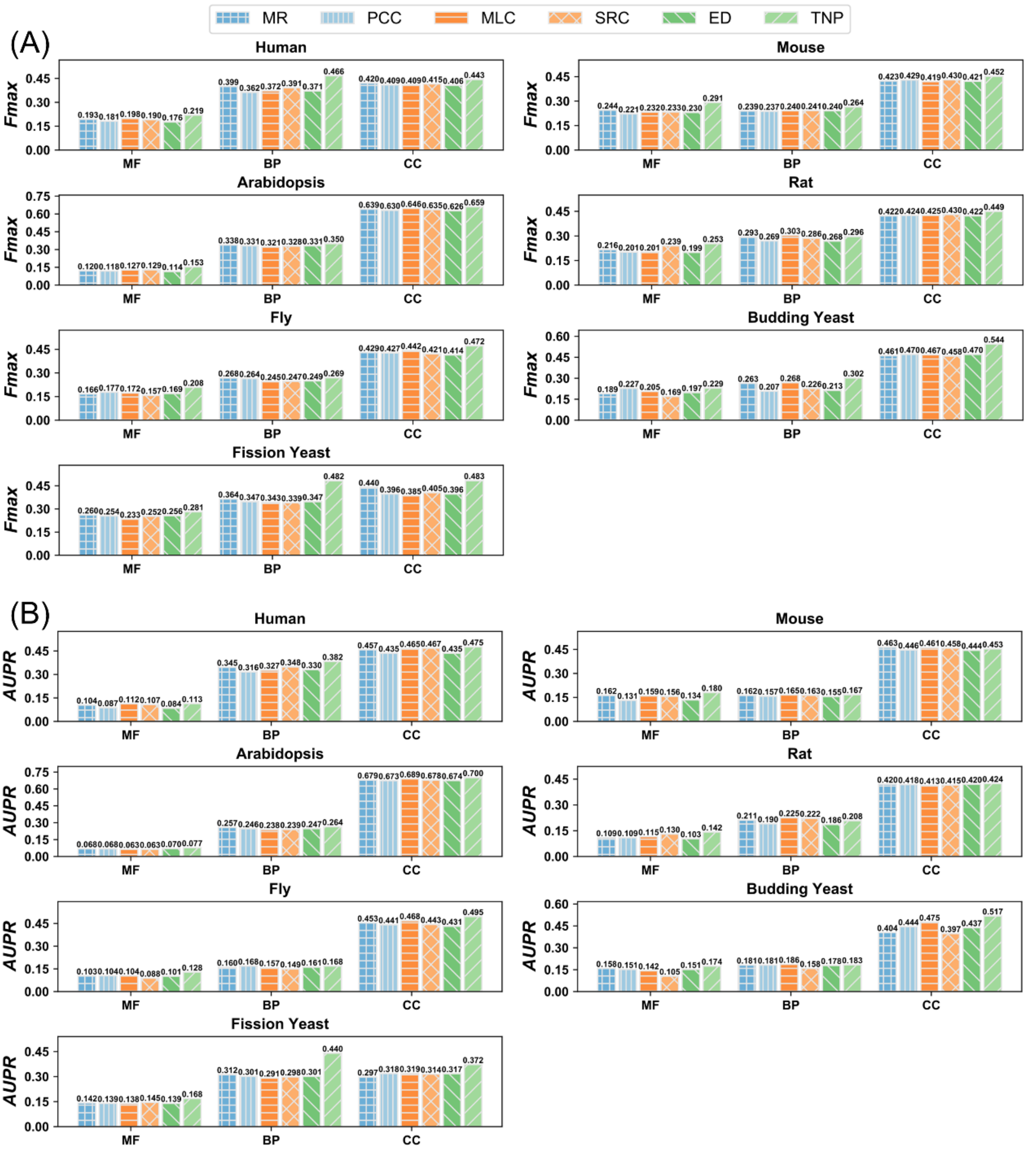


**Figure S12 The performance of six expression profile-based methods for 7 species on CAFA3 test dataset**

**A**. The $Fmax$ values of six methods. **B**. The $AUPR$ values of six methods.

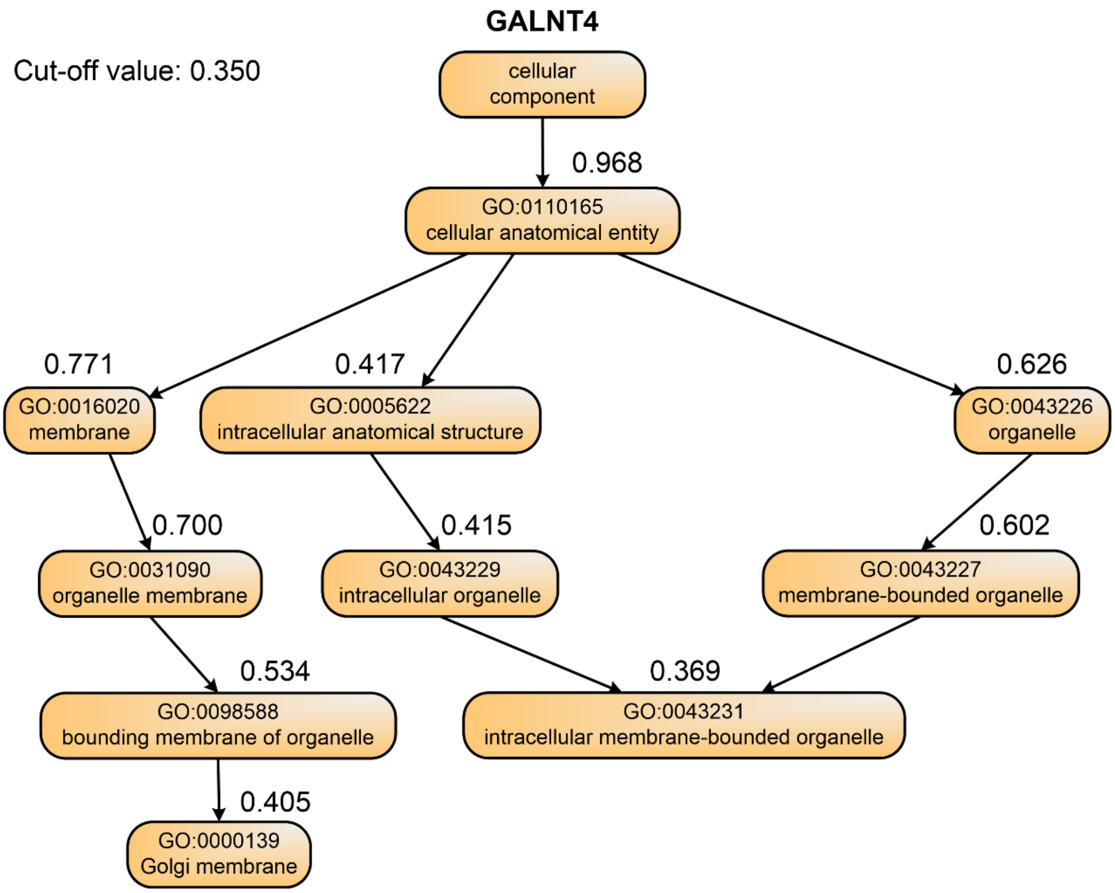


**Figure S13 The directed acyclic graph of predicted GO terms with corresponding confidence scores for gene GALNT4 by TripletGO.**

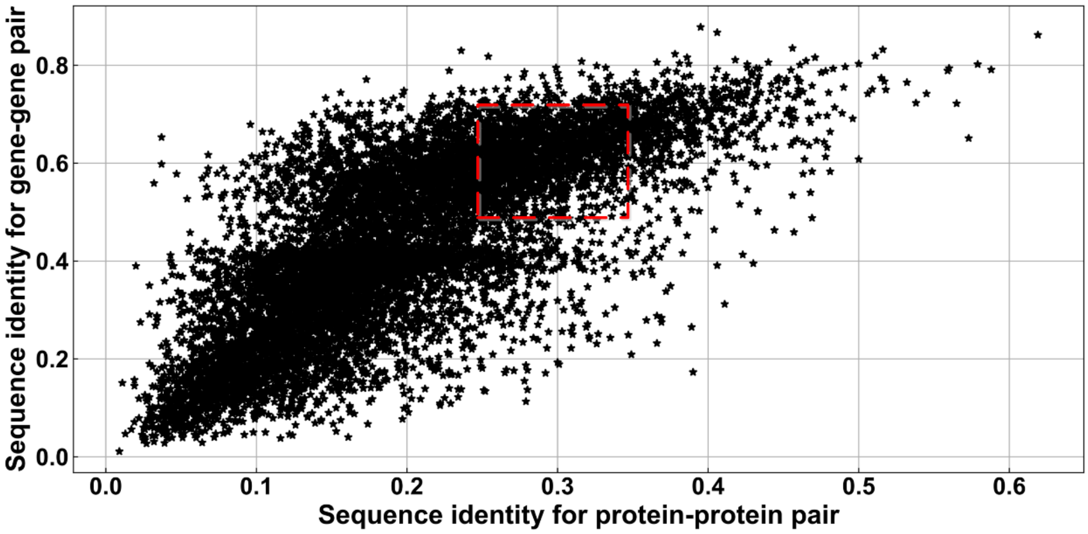


**Figure S14 The distribution of sequence identities for 10000 gene-gene pairs and 10000 mapped protein-protein pairs**
